# Metastatic niche mediated activation of metastasis initiating cells in ovarian cancer through miR-193b-3p downregulation via the ERK/EZH2/DNMT1 axis

**DOI:** 10.1101/2025.06.17.660246

**Authors:** Subramanyam Dasari, Ji Wang, Frank Cheng, Melissa Halprin, David Pepin, Yang Yang-Hartwich, Anirban K Mitra

## Abstract

Extensive metastasis at the time of diagnosis is a major contributor to the poor prognosis of ovarian cancer (OC) patients. There is a critical need to better understand the mechanism of regulation of metastasis to develop effective treatment strategies targeting the process. Metastasis initiating cells (MICs) have cancer stem cell-like properties along with the ability to invade. Their potential role in OC is unique as the dissemination from the primary tumors involves passive processes like exfoliation. However, the role of MICs during OC metastatic colonization is critical and poorly understood. Using an organotypic 3D culture model of the human omentum, we have studied the productive crosstalk between OC MICs and the metastatic microenvironment. We report the role of miR-193b-3p, a clinically relevant metastasis suppressor microRNA, which is downregulated in the OC by paracrine signals from the microenvironment, inducing the MIC phenotype. Using heterotypic coculture models, conditioned medium experiments, secretome analysis, inhibition, and rescue experiments, we show that bFGF and IGFBP6 secreted by mesothelial cells in the microenvironment induce miR-193b-3p downregulation in OC MICs via the ERK/EZH2/DNMT1 axis. The miR-193b-3p downregulation induced an increased expression of its target cyclin D1, which imparted a cancer stem cell phenotype. Urokinase, another target of miR-193b-3p, induced invasive growth. Together, these targets impart the MIC phenotype to the OC cells. miR-193b-3p replacement therapy could suppress metastasis in a patient derived xenograft model of OC metastasis, indicating the translational potential of this approach to target MICs in OC patients.

## Introduction

Ovarian cancer (OC) is the deadliest gynecological malignancy with a 5-year survival rate of <50% and high-grade serous OC (HGSOC) is the most prevalent and lethal subtype^1,2^. A key reason for this is that most HGSOC patients present with disseminated disease at the time of diagnosis ^2^. OC predominantly undergoes transcoelomic metastasis with the cancer cells exfoliating from the primary tumor into the peritoneal fluid. Subsequently, they are dispersed through the peritoneal cavity and attach to the organs therein. A common site is the omentum, a fat pad formed by the double fold of the peritoneal membrane. To form metastases, the OC cells must first attach to the surface of the omentum, invade through the mesothelium covering its surface and the underlying basement membrane. Here, they encounter a microenvironment that is very different from the one they had experienced in the primary tumor. Therefore, only those cells that can adapt to this new microenvironment will successfully colonize the site of metastasis. The process of adaptation involves productive reciprocal interactions with the metastatic microenvironment, including mesothelial cells, fibroblasts, extracellular matrix (ECM), and other stromal cells, causing changes in gene expression that promote metastatic colonization.

Recent studies have revealed the existence a subpopulation of cancer cells, called metastasis initiating cells (MICs), which are very plastic in nature and have the capacity to establish metastases ^3,4^. MICs are characterized by a cancer stem cell like phenotype, ability to productively interact with the metastatic niche, and adapt to external stresses enabling them to seed the metastases, also potentially cause relapse ^3,4^. Our understanding of the mechanisms regulating the MIC phenotype and their interactions with the metastatic niche in OC is limited.

microRNAs are a class of small, non-coding RNAs that regulate a wide range of biological process through sequence specific translational inhibition of their target genes. We have previously used an organotypic 3D culture system mimicking the surface layers of the omentum to study the mechanism of regulation of early OC metastatic colonization by microenvironment regulated microRNAs^5^. Later, we identified microRNAs that are deregulated in early and advanced metastases using a combination of the 3D omentum culture model matched primary tumors and metastases from HGSOC patients respectively ^6^. Using these approaches, we have identified miR-193b-3p as a key microRNA, which is decreased in the metastasizing OC cells through their interactions with the metastatic niche. miR-193b-3p down regulation decreased metastasis *in vitro, ex vivo,* and *in vivo* ^5^. It is not well established that the weather endogenous miRNAs are involved in the regulation of metastatic initiation as well as recurrence of diseases. miRNAs have been shown to reprogram somatic cells into pluripotent stem cells^7^ and regulate OC stem cells^8^.

In the present study, we demonstrate the regulation of OC MICs by the downregulation of miR-193b-3p. We report the mechanism by which paracrine signals from the metastatic niche downregulated miR-193b-3p in OC cells via the ERK/EZH2/DMNT1 axis. Finally, we have identified Cyclin D1 as the downstream effector of miR-193b-3p, which helps induce the MIC phenotype. Our data offer novel insights into the regulation of MICs by the crosstalk between the metastatic niche and OC cells, thus laying the foundations for the development of miR-193b-3p replacement therapy to target OC MICs.

## Results

### Metastasis initiation and induction of stemness by miR-193b-3p

Since MICs have been reported to have stem cell like characteristics, we started by testing the ability of OC stem cells to attach to the omentum and invade through its outer layers. Aldehyde dehydrogenase (ALDH) activity is a well-established marker for OC stem cells^9–11^. OVCAR3 HGSOC cells were sorted into ALDH+ and ALDH-cells following ALDEFLUOR assay. *In vivo* adhesion assay was performed by injecting them i.p. in female NSG mice. ALDH+ OC stem cells had significantly higher adhesion to the mouse omentum than the ALDH-cells (Figure 1A, Suppl Figure 1A). The ability of the OVCAR3 cells to invade through the surface layers of the omentum was assayed by seeding fluorescent OVCAR3 cells on the 3D omentum culture assembled in FluoroBlok transwell inserts and measuring the number of fluorescent cells that invade to the bottom of the insert. Again, ALDH+ OVCAR3 stem cells had significantly higher capacity to invade through the outer layers of the omentum (Figure 1B, Suppl Figure 1B). Since we had previously demonstrated that miR-193b-3p downregulation was important for OC metastasis initiation, we tested the levels of miR-193b-3p in OC stem cells. ALDH+ OC cells had significantly lower miR-193b-3p expression (Figure 1C). OC spheroids grown in ultra-low adhesion plates are enriched for OC stem cells and had low expression of miR-193b-3p (Figure 1C, Suppl Figure 2A). Low expression of miR-193b-3p results in poor survival in OC patients (Suppl Fig 2B). Since we have previously shown that signals from the mesothelial cells in the metastatic microenvironment induce miR-193b-3p downregulation^5^, we tested the effect of mesothelial cell conditioned medium on the expression of a panel of stem cell markers in OC cells. Mesothelial cell conditioned medium could induce the expression of ALDH1A1, OCT4, NANOG, and SOX2 in all 3 OC cells (Figure 1D). Conversely, overexpression of miR-193b-3p caused a decrease in the expression of these cancer stem cell markers (Figure 1E). Similarly, miR-193b-3p overexpression decreased ALDH activity and spheroid formation in ultra-low adhesion plates (Figure 1F and G). Taken together, our data indicates that OC stem cells are more capable of attaching to and invading through the outer layers of the omentum – key steps in the initiation of metastatic colonization. Moreover, miR-193b-3p downregulation, which is important for initiation of colonization, is associated with the OC stem cell phenotype.

**Figure 1.**
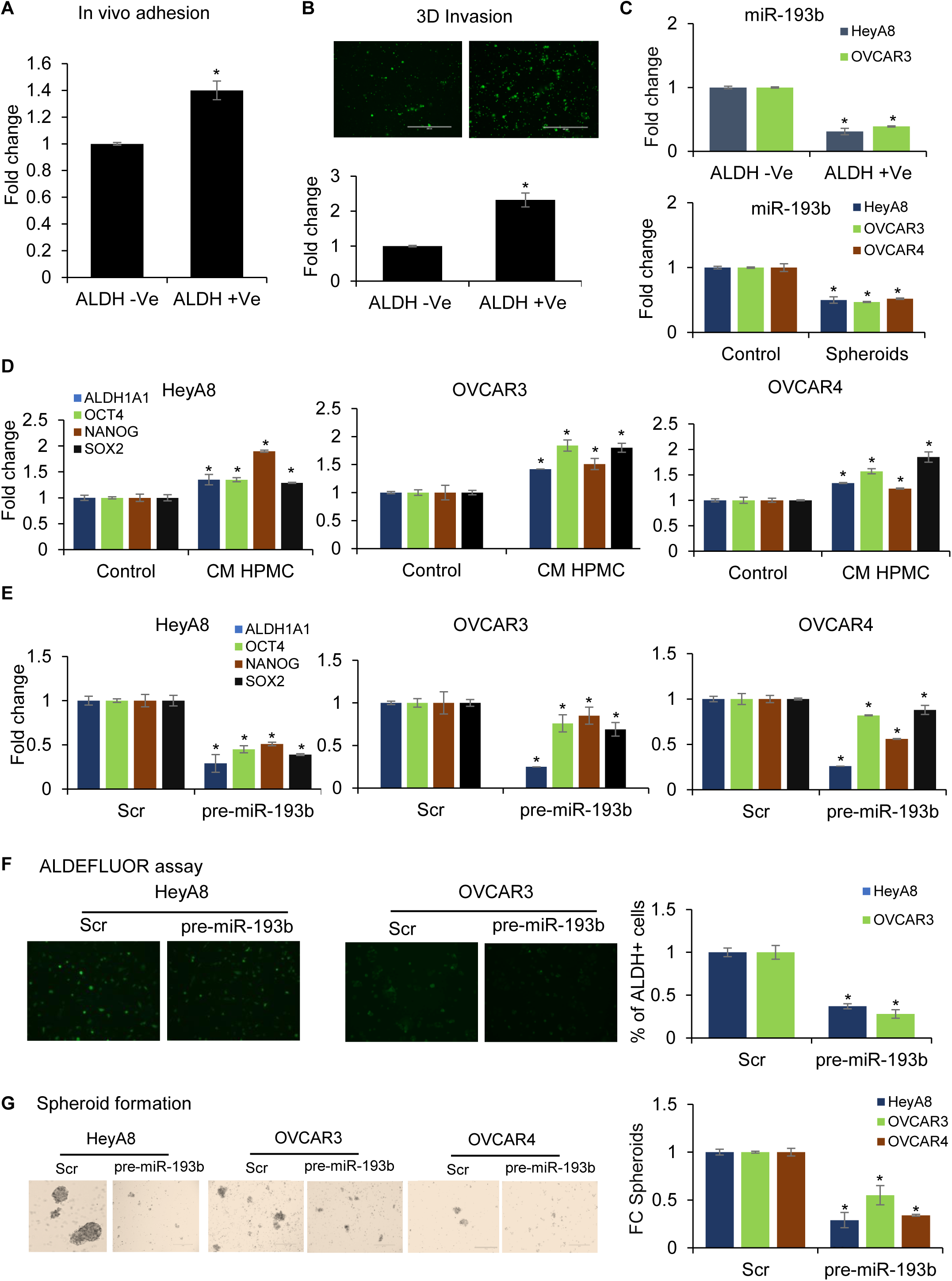
MICs and miR-193b-3p. **(A) 3D invasion:** OVCAR3 cancer stem cells with high aldehyde dehydrogenase (ALDH) activity, were separated by ALDEFLUOR assay and fluorescence activated cell sorting (FACS). The ALDH +ve and -ve cells were labelled with CMFDA and seeded on the 3D omental culture that was assembled in FluoroBlok transwell inserts. The OC cells were allowed to invade through the 3D omental culture for 16 h, imaged (5 fields/insert) using the EVOS FL auto microscope, and quantified. DMEM with 10% FBS served as the chemoattractant in the lower chamber. **(B). In vivo adhesion:** NSG mice were injected i.p. with CMFDA labeled ALDH +ve or ALDH -ve OVCAR3 cells (350K/0.5 mL/mouse, 3 mice per group). After 3 hours, mice were euthanized to collect their omentum and peritoneum with adherent OC cells. They were washed and the adherent cells isolated with trypsin and the fluorescence quantified using a SynergyH1 plate reader. **(C) *Top:* miR-193b-3p in ALDH +Ve and -Ve cells:** OC stem cells (HeyA8, OVCAR3) were isolated using ALDEFLUOR assay and FACS. RNA was isolated, from both ALDH +ve and -ve OC cells and qRT-PCR was performed for miR-193b-3p. ***Bottom:* miR-193b-3p in 2D vs Spheroids:** OC cells (HeyA8, OVCAR3 and OVCAR4) were grown in a 2D culture plates and ultra-low-adherent plates for 4-10 days for spheroid formation. RNA was isolated and qRT-PCR was performed for miR-193b-3p. **(D). Stem cell markers induced by CM:** OC cells (HeyA8, OVCAR3 and OVCAR4) cells were grown in conditioned medium (CM) collected from human primary mesothelial cells (HPMC) while controls were grown in serum free medium. qRT-PCR was done to test the effect on expression of stem cell markers. **(E). Overexpression of miR-193b-3p:** HeyA8, OVCAR3 and OVCAR4 cells were transfected with pre-miR-193b-3p or scrambled control oligos (Scr) and used for evaluating cancer stemness. RNA was isolated, and qRT-PCR was performed for stem cell markers. **(F) ALDEFLUOR Assay:** ALDH enzymatic activity was measured using ALDEFLUOR assay. ***(Left)*** Representative fluorescent image of ALDH +ve cells (green) taken using the EVOS FL auto microscope. ***(Right)*** ALDH +ve cells population was quantified by LSRII flow cytometer (BD Biosciences) and the average number of ALDH +ve were plotted. **(G) Spheroid formation:** Transfected cells were grown in an ultra-low-attachment 24-well plates for a period of 4-10 days for spheroid formation. Spheroids were imaged with EVOS FL Auto microscope, counted, and the fold change (FC) in spheroids was plotted. All error bars represent mean ± SD; 3 independents experiments, * p<0.01, Students t-test.

### Downregulation of miR-193b-3p by mesothelial cells

We have previously demonstrated that the miR-193b-3p downregulation in OC cells by mesothelial cells in the metastatic microenvironment is important for metastatic colonization^5^. This was through induction of DNMT1 by the mesothelial cells, which caused miR-193b-3p promoter hypermethylation^5^. However, the underlying mechanism was not clear. Therefore, the first thing we tested was if the signals from the mesothelial cells were of a paracrine or juxtacrine nature. We have developed a proximal culture assay specifically to address this question^12^. This assay differentiates between paracrine and juxtacrine signaling by seeding cells on either surface of a 10 µm thick membrane with 0.4 µm pores, which allows exchange of secreted factors between cells at the localized high concentrations while not permitting direct contact (Supplementary Figure 3A). Proximal culture of OC cells with mesothelial cells resulted in decreased expression of miR-193b-3p in the OC cells, suggesting a paracrine mechanism (Figure 2A, left). To rule out the possibility of exchange of signals via tunneling nanotubules, we tested the effect of concentrated conditioned medium form mesothelial cells on miR-193b-3p expression of OC cells. Again, secreted factors from mesothelial cells downregulated miR-193b-3p in OC cells (Figure 2A, right). DNMT1 is the only DNMT isoform induced in the OC cells by the metastatic microenvironment^5^. We confirmed that coculture with mesothelial cells could induce DNMT1 in a panel of OC cells (Figure 2B, left) and this was through a paracrine signaling mechanism, as demonstrated by proximal culture with mesothelial cells and conditioned medium experiments (Figure 2B, middle and right). Treatment with a DNMT inhibitor (decitabine) resulted in an increased expression of miR-193b-3p in a panel of OC cells, confirming the role of DNMT in miR-193b-3p downregulation (Figure 2C). miR-193b-3p has been reported to be downregulated by ERK in pancreatic cancer^13^ and DNMT1 can potentially be regulated by ERK in gastric cancer^14^. Therefore, we tested the effect of ERK inhibition on miR-193b-3p and DNMT1 expression in a panel of OC cells (Figure 2D). ERK inhibition increased miR193b and decreased DNMT1 expression in all OC cells, indicating its role in the pathway activated by mesothelial cells that leads to miR-193b-3p downregulation. Next, we tested the effect of the metastatic microenvironment on ERK activation and DNMT1 induction in OC cells by immunofluorescence using the 3D omentum culture (schematic in Supplementary Figure 3B). ERK1/2 phosphorylation was induced in the OC cells seeded on the 3D omentum culture while there was no change in the total ERK1/2 expression (Figure 2E). The 3D omentum culture also induced DNMT1 expression in the OC cells (Figure 2E). Similarly, ERK1/2 phosphorylation and DNMT1 protein expression was induced in OC cells treated with mesothelial cell conditioned medium, confirming the involvement of a paracrine mechanism in their induction (Figure 2F).

**Figure 2.**
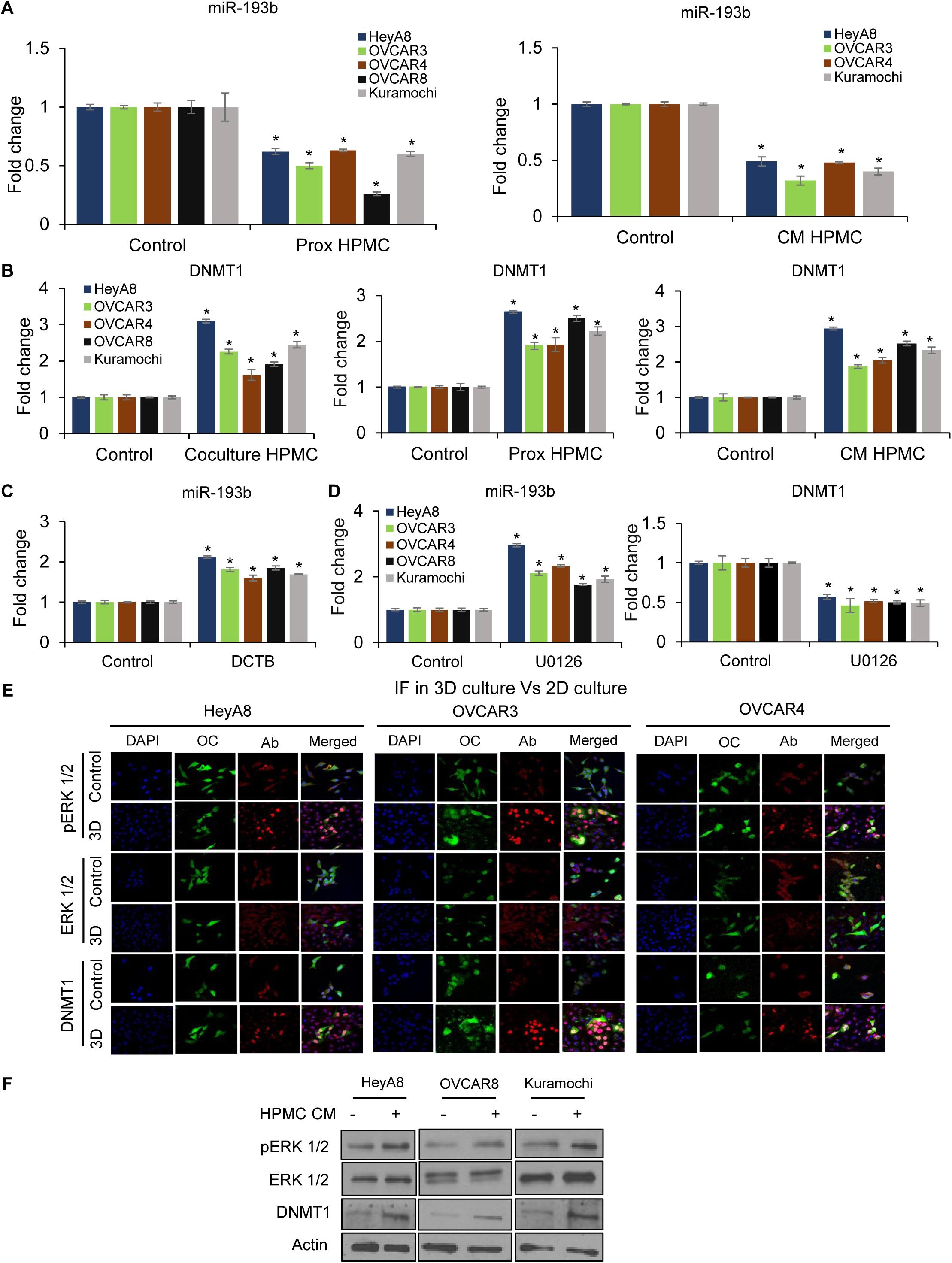
Paracrine signals from mesothelial cells downregulate miR-193b-3p: **(A). Proximal culture *(Left)*:** OC cells (HeyA8, OVCAR3, OVCAR4, OVCAR8 and Kuramochi) were grown in a proximal culture setup with HPMC for 48 hours as described in methods. RNA isolated and qRT-PCR performed for miR-193b-3p expression compared corresponding controls, which did not have the HPMC seeded on the opposite surface. **Conditioned medium *(Right)*:** OC cells were grown in condition medium from HPMC were used for RNA isolation and qRT-PCR for miR-193b-3p expression compared to corresponding controls, grown in serum free medium. **Mesothelial signals induce DNMT1 expression (B) Coculture *(left)*:** GFP-labeled OC cells were cocultured with mesothelial cells (HPMC). After 2 days, the OC cells were isolated using FACS and used for RNA isolation and qRT-PCR for miR-193b-3p expression compared to corresponding monoculture controls. **Proximal culture *(middle)*:** OC cells were grown in a proximal culture setup with HPMC, for 48 hours. RNA isolated and qRT-PCR performed for DNMT1 expression compared corresponding controls, which did not have the HPMC seeded on the opposite surface. **Conditioned medium *(Right)*:** OC cells were grown in conditioned medium from HPMC were used for RNA isolation and qRT-PCR for DNMT1 expression compared to corresponding controls, grown in serum free medium. **(C) DNMT inhibition:** OC cells (HeyA8, OVCAR3, OVCAR4, OVCAR8 and Kuramochi) were treated with DCTB (5 µM, daily). RNA was isolated after 4 days of treatment and the expression of miR-193b-3p was compared with corresponding vehicle controls, using qRT-PCR. **(D) ERK inhibition:** OC cells (HeyA8, OVCAR3, OVCAR4, OVCAR8 and Kuramochi) were treated with U0126 (10 µM). RNA isolated after 48 h of treatment, and the expression of miR-193b-3p (left) and DNMT1 (right) was compared with corresponding vehicle controls, using qRT-PCR. **(E) Immunostaining on 3D omentum culture:** GFP-labeled OC cell (HeyA8, OVCAR3 and OVCAR4) were seeded on the 3D omentum culture and allowed to incubate for 24 h, fixed and stained for phosphorylated ERK1/2 (red), total ERK1/2 (red), and DNMT1 (red). Nuclei were stained with Hoechst 33342 (blue), allowing visualization of the nonfluorescent constituents of the 3D omentum culture. **(F)** Immunoblots for phosphorylated ERK1/2, ERK1/2, and DNMT1 in OC cells treated with mesothelial cell conditioned medium (HPMC CM). Representative image shown from 3 independent experiments. All error bars represent mean ± SD; 3 independents experiments, * p<0.01, Students t-test.

The next question was to identify the link between ERK activation and recruitment of DNMT1 to miR-193b-3p promoter. ERK has been reported to regulate EZH2 expression in lung and breast cancer^15,16^. Also, histone H3 lysine 27 trimethylation (H3K27me3) induced by EZH2 recruits DNMT1 to catalyze promoter hypermethylation^17^. Therefore, we tested the potential role of EZH2 in regulating miR-193b-3p and OC stem cells. Treatment with an EZH2 inhibitor (GSK126) increased miR-193b-3p expression and decreased ALDH1A1 expression in OC cells (Figure 3A and B). Interestingly, ERK inhibition led to decreased levels of DNMT1, EZH2, and H3K27me3 in a panel of OC cells, indicating that EZH2 and DNMT1 are regulated by ERK in OC cells (Figure 3C). Next, we wanted to test if EZH2 was indeed induced in OC cells by the metastatic microenvironment. Therefore, immunofluorescence staining for EZH2 was performed in GFP expressing OC cells seeded on the 3D omentum culture and compared to 2D control (Figure 3D). EZH2 expression was induced in OC cells by the 3D culture, further confirming its induction by the metastatic microenvironment. Finally, we tested the localization of EZH2, H3K27me3, and DNMT1 at the miR-193b-3p promoter by ChIP-qPCR. Treatment with ERK inhibitor decreased their localization to the miR-193b-3p promoter (Figure 3E). MeDIP-qPCR was done to test for miR-193b-3p promoter methylation. Again, ERK inhibition decreased miR-193b-3p promoter methylation (Figure 3E). Taken together, our data indicates that paracrine signals form mesothelial cells in the metastatic microenvironment activated ERK in the OC cells, which led to EZH2 and DNMT1 induction. EZH2 recruitment to the miR-193b-3p promoter caused H3K27me3 at the miR-193b-3p promoter, recruiting DNMT1 that catalyzed miR-193b-3p promoter hypermethylation, decreasing its expression.

**Figure 3.**
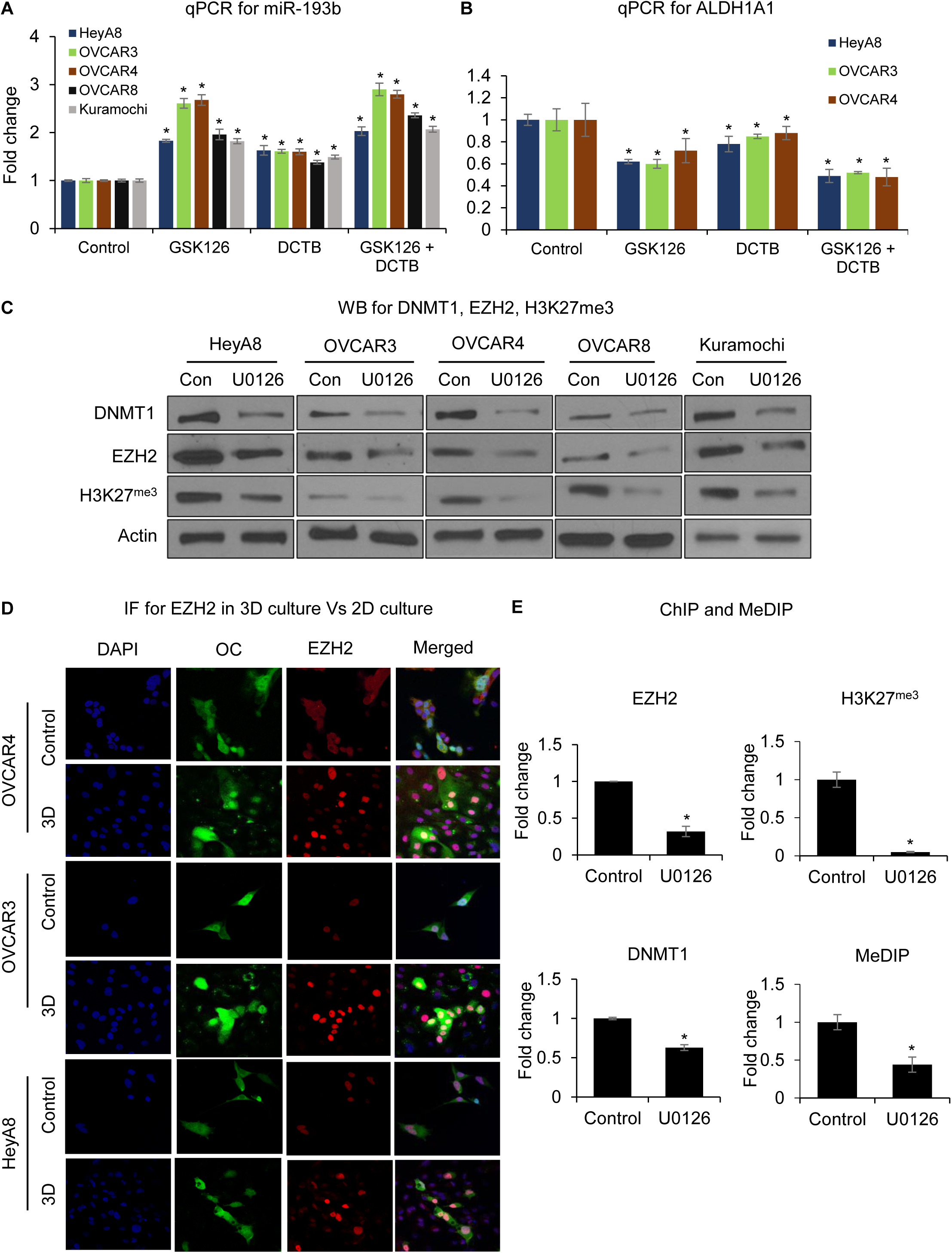
Regulation of miR-193b-3p. OC cells (HeyA8, OVCAR3, OVCAR4, OVCAR8 and Kuramochi) were treated with 10 µM GSK 126, 5 µM decitabine (DCTB) or a combination of both daily. RNA was isolated after 4 days followed by qRT-PCR for the expression of miR-193b-3p **(A)** and the OC stem cell marker ALDH1A1 (**B),** compared to DMSO controls. **(C)** OC cells (HeyA8, OVCAR3, OVCAR4, OVCAR8 and Kuramochi) were treated with U0126 (10 µM) or DMSO (control). Protein was isolated after 48 h of treatment for immunoblotting for DNMT1, EZH2 and H3K27me3. Representative image shown from 3 independent experiments. **(D) Immunofluorescence on 3D culture:** GFP-labeled OC cell (HeyA8, OVCAR3 and OVCAR4) were seeded on the 3D omentum culture, allowed to incubate for 24 h, and then fixed and stained for EZH2 (red). Nuclei were stained with Hoechst 33342 (blue), allowing visualization of the nonfluorescent constituents of the 3D omentum culture. **(E)** OC cells (HeyA8) were treated with U0126 (10 µM) and used to perform ChIP. Cells were cross-linked with formaldehyde, chromatin extracted, sheared and used for pulldown with specific antibodies. Cross-linked DNA was released from the antibody-captured protein-DNA complex and purified with Fast-Spin columns. Eluted DNA was used for assessing the enrichment of EZH2, H3K27^Me3^ and DNMT1 at miR-193b-3p promoter using qRT-PCR. Alternately, methylated DNA Immunoprecipitation (MeDIP) was performed with the U0126 treated HeyA8 cells. Single-stranded, fragmented genomic DNA containing 5-methylcytosine was specifically captured with a monoclonal 5-mC antibody in the presence of a bridging antibody and magnetic beads. The eluted DNA was used to analyze the enrichment of 5-methylcytosine (5-mC) at the miR-193b-3p promoter using qRT-PCR. All error bars represent mean ± SD; 3 independents experiments, * p<0.01, Students t-test.

### miR-193b-3p induces OC stem cells through increased expression of its target, CCND1

Having determined the mechanism of downregulation of miR-193b-3p, the next step was to determine its mechanism by which its downregulation induces stemness. Since microRNAs act through the downregulation of their targets, RNA-seq was performed to compare gene expression changes in OVCAR3, OVCAR4, and HeyA8 OC cells transfected with pre-miR-193b-3p or scrambled negative control. Only the 677 genes commonly downregulated in all 3 cells overexpressing miR-193b-3p were considered for subsequent analysis, to avoid cell specific effects (Figure 4A). Using microRNA target prediction software Targetscan, miRmap and miRwalk, these 677 genes were analyzed for predicted miR-193b-3p seed matches. 43 genes were predicted miR-193b-3p targets by at least two software. A pathway analysis of these 43 genes using Metascape^18^ showed 5 key pathways, which were all involved in differentiation and development (Figure 4B). Cyclin D1 (CCND1) was selected for functional testing based on a literature search for cancer stemness relevant genes among these potential targets^19,20^. We have previously identified Urokinase-type Plasminogen Activator (uPA) as a direct target of miR-193b-3p, regulating invasiveness and proliferation during metastasis^5^. Both CCND1 and uPA mRNA were found to be downregulated in a panel of OC cells, when miR-193b-3p was overexpressed in them (Figure 4C). The inverse relationship between miR-193b-3p and its targets was further tested by western blotting following transfection of OC cells with pre-miR-193b-3p or anti-miR-193b-3p (Figure 4D). Both cyclin D1 and uPA were downregulated upon miR-193b-3p overexpression and upregulated upon miR-193b-3p inhibition. We have previously demonstrated the role of uPA in OC metastasis^5^. Next, the role of Cyclin D1 as a functional effector of miR-193b-3p on OC stem cell induction was studied. Knocking down cyclin D1 resulted in a decrease in OC stem cell markers, ALDH activity, and spheroid formation in ultra-low adhesion plates, while overexpression of cyclin D1 increased spheroid formation (Figure 4 E-G and Suppl Figure 4A-C). Although this confirmed the role of cyclin D1 in regulating OC stem cell function, it does not necessarily implicate it as a downstream effector if miR-193b-3p. Therefore, a functional rescue experiment was performed to test its role as a downstream effector of miR-193b-3p. A panel of OC cells were transfected with pre-miR-193b-3p and cyclin D1 expression vector or their respective controls, scrambled negative control oligo and empty vector (Suppl Figure 5), to test the effect of cyclin D1 on stemness suppression by miR-193b-3p overexpression. Cyclin D1 overexpression was found to rescue the ALDH1A1 downregulation induced by miR-193b-3p (Figure 5A). Similarly, Cyclin D1 overexpression rescued miR-193b-3p induced suppression of spheroid formation in the OC cells (Figure 5B). Taken together, the data confirmed that cyclin D1 was the downstream effector of miR-193b-3p in the regulation of OC stem cells.

**Figure 4.**
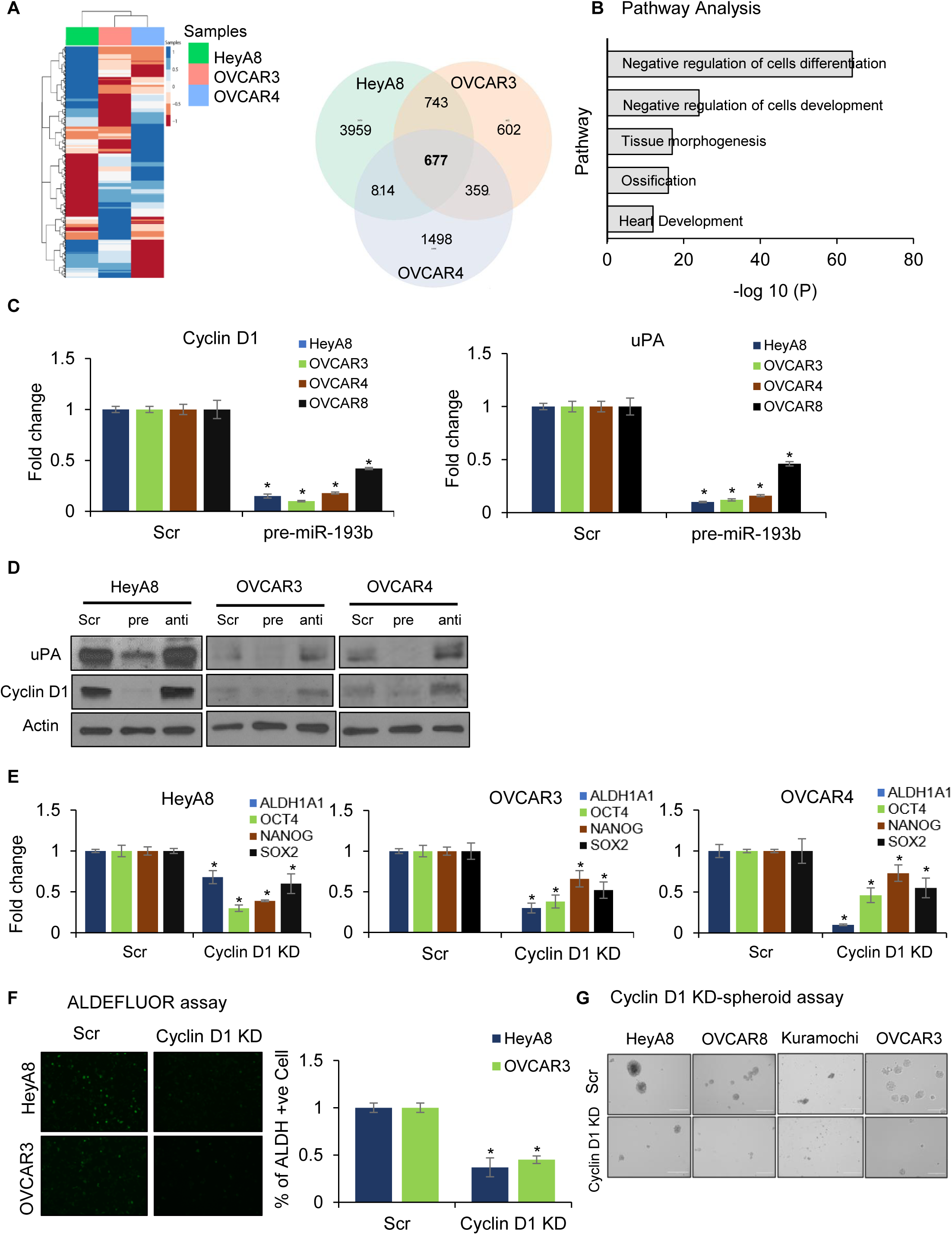
miR-193b-3p target identification. **(A)** RNA-seq was done using HeyA8, OVCAR3 and OVCAR4 cells transiently transfected with pre-miR-193b-3p or scrambled control oligos (n=3/group). ***Left:*** Heat map representing the deregulated (both up and down regulated) genes in response to miR-193b-3p overexpressing HeyA8, OVCAR3 and OVCAR4 cells compared to the corresponding controls. ***Right:*** Venn diagram showing the number of genes downregulated uniquely and commonly in all the three OC, cells upon overexpression of miR-193b-3p, compared to the respective controls. **(B)** Pathway analysis of the predicted targets of miR-193b-3p among the 677 commonly downregulated genes was performed using Metascape. **(C) Target validation by qRT-PCR:** qRT-PCR for CCND1 and uPA done using RNA from HeyA8, OVCAR3 and OVCAR4 cells transfected with pre-miR-193b-3p or scrambled control oligos (scr). **(D) Target validation by Immunoblotting:** HeyA8, OVCAR3 and OVCAR4 cells were transfected with pre-miR-193b-3p or anti-miR-193b-3p or scrambled control oligos (scr) and cells were lysed after 48 h. Immunoblotting for CCND1 and uPA was performed. Representative image shown from 3 independent experiments. (**E) Functional role of cyclin D1:** HeyA8, OVCAR3 and OVCAR4 cells were transfected with CCND1 siRNA or scrambled control oligos (scr) and cells were lysed after 48 h. qRT-PCR was performed for OC stem cell markers (ALDH1A1, OCT4, NANOG and SOX2). **(F) ALDEFLUOR assay:** ALDEFLUOR assay performed with OC cells (HeyA8 and OVCAR3) transfected with CCND1 siRNA or scrambled control oligos (scr). Fluorescent images of ALDEFLUOR +ve cells were taken using an EVOS FL auto microscope (***left***). Similarly, ALDH +ve cell population was quantified by LSRII flow cytometer (BD Biosciences) and the average number of ALDH +ve cells was plotted ***(right)***. **(G) Spheroid formation assay:** Spheroid formation assay was performed with OC cells transfected with CCND1 siRNA or scrambled control oligos (scr). Images of spheroids were taken was taken using an EVOS FL auto microscope. Representative images from 3 independent experiments are shown. All error bars represent mean ± SD; 3 independents experiments, * p<0.01, Students t-test.

**Figure 5:**
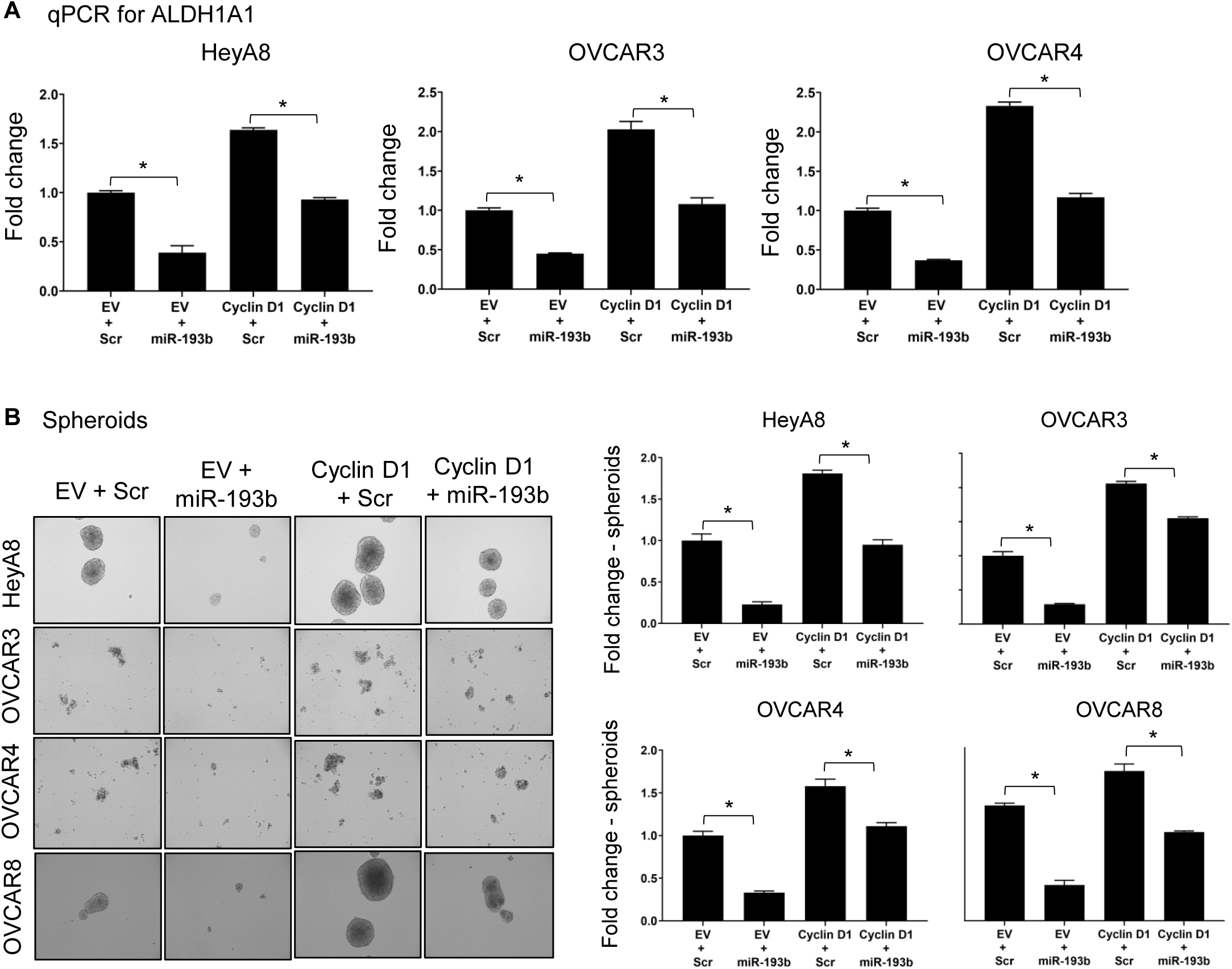
Cyclin D1 is a functional effector of miR-193b-3p: **(A)** OC cells (HeyA8, OVCAR3, OVCAR4 and OVCAR8) were co-transfected with pre-miR-193b-3p and CCND1 plasmid and RNA isolated after 48 h. qRT-PCR was performed for the expression of OC stem cell marker ALDH1A1. **(B)** OC cells (HeyA8, OVCAR3, OVCAR4 and OVCAR8) were co-transfected with different combinations of pre-miR-193b-3p, cyclin D1 expression plasmid, scrambled control oligo (Scr), and empty vector (EV). Cells were seeded in ultralow-adhesion plates for spheroid formation for a period of 4-10 days. Representative images of spheroids from 3 independent experiments are shown and quantification of the fold change in spheroids was plotted. All error bars represent mean ± SD; 3 independents experiments, * p<0.01, Students t-test.

### Regulation of miR-193b-3p by microenvironmental signals

Having demonstrated cyclin D1 as the miR-193b-3p effector, the next step was to identify the microenvironmental signals that induce miR-193b-3p downregulation in metastasizing OC cells. Secreted factors from mesothelial cells were found to downregulate miR-193b-3p (Figure 2A). Therefore, to identify the secreted factors unique to mesothelial cells, a secretome array was employed. Conditioned medium from mesothelial cells, OC cells or OC cells cocultured with mesothelial cells were analyzed using secretome arrays (Figure 6A and suppl figure 6A). Basic fibroblast growth factor (bFGF) and insulin like growth factor binding protein 6 (IGFBP6) were found to be higher in mesothelial cell secretome compared to OC cells. Interestingly, bFGF was further induced in the mesothelial cell OC coculture. To test the clinical relevance of these findings, we analyzed scRNA-seq data from 11 OC patient metastases^21^. Both bFGF and IGFBP6 are predominantly expressed by stromal cells with minimal expression in the cancer cells and low expression in some subpopulations of immune cells (Figure 6B). The stromal and cancer components were further resolved into subpopulations of mesothelial cells (Meso 0-1), endothelial cells (Endo 0-1), cancer associated fibroblasts (CAFs) (FB 0-8), and cancer cells (CC 0-12) and analyzed for their expression of bFGF and IGFBP6 (Figure 6C). While mesothelial cells, endothelial cells, and CAFs had higher expression of bFGF and IGFBP6 than the cancer cells, there was heterogeneity in their expression within each cell type. This confirmed that the stromal cells in patient metastasis were the dominant source of bFGF and IGFBP6 and CAF and mesothelial cells were key contributors. However, during the initial steps of metastatic colonization, mesothelial cells serve as the main source of these secreted factors as there are no CAFs at that stage. The expression of FGF2 and IGFBP6 mRNA was further confirmed to be higher in mesothelial cells and CAFs as compared to OC cells (Suppl Figure 6B).

**Figure 6:**
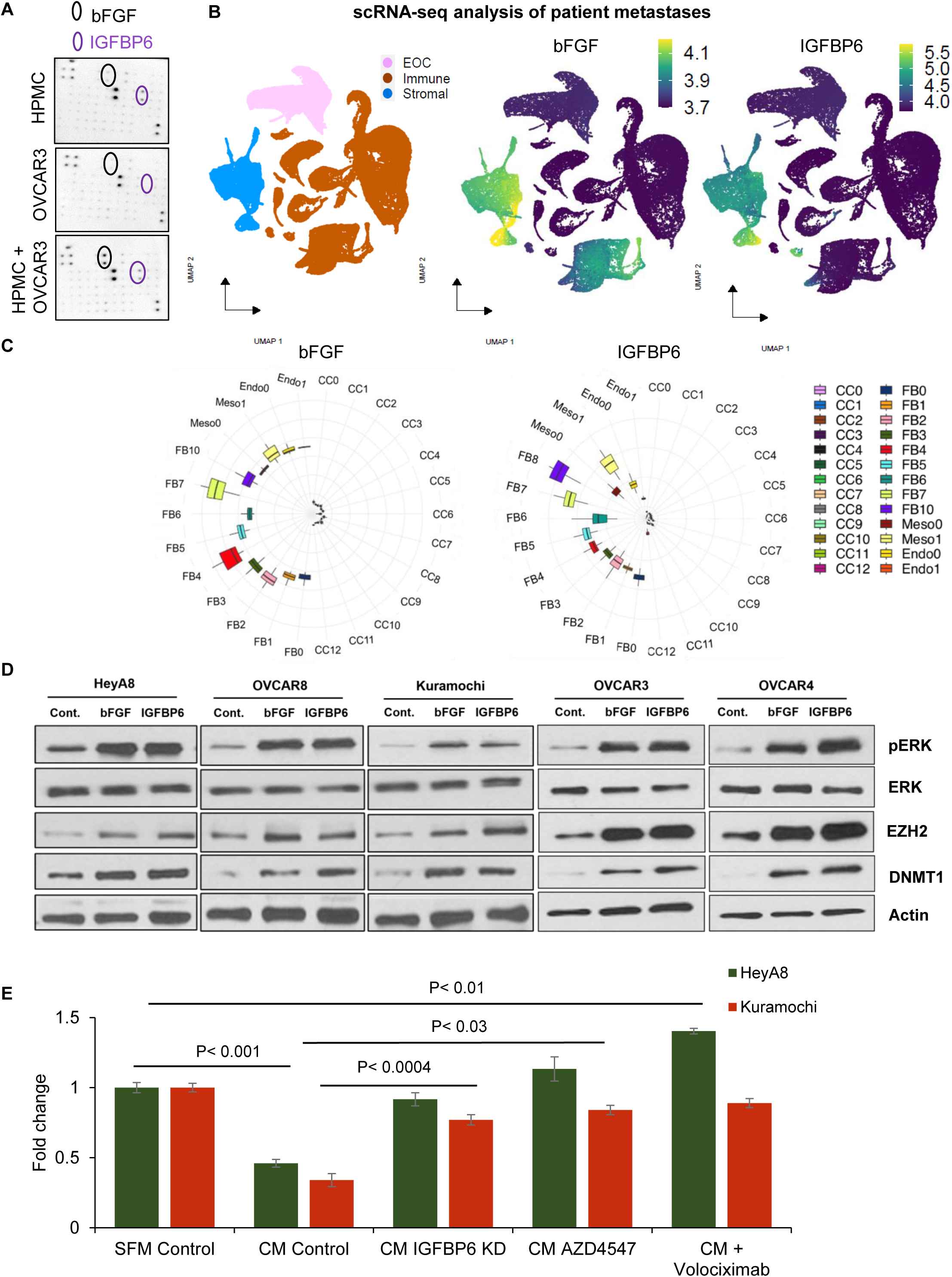
Crosstalk with the microenvironment: **(A)** Secretome array: Conditioned medium was collected from human primary mesothelial cells, OVCAR3 cells and, their coculture, to screen the secreted factors using the Human Growth Factor Antibody Array C1, (RayBiotech, Inc.). Representative image of the array with the bFGF (black) and IGFBP6 (purple) spots encircled. **(B)** Analysis of scRNA-seq from metastatic tumors from 11 OC patients was analyzed and segregated into OC (EOC), stromal, and immune cell subpopulations (UMAP on left). UMAPs depicting the expression of bFGF (middle) and IGFBP6 (right) in these subpopulations. Bright yellow indicates high expression while dark purple denotes low expression. **(C)** The stromal (Endo 0-1, Meso 0-1, FB 0-10) and OC (CC 0-12) cell populations were further segregated into subpopulations and the expression of bFGF and IGFBP6 were analyzed in these subpopulations. Each Circos plot shows the expression levels of the respective genes in each subpopulation as box and whisker plots. **(D)** OC cell (HeyA8, OVCAR3, OVCAR4, OVCAR8, and Kuramochi) were treated with recombinant human bFGF (1.5 ng/mL) or IGFBP6 (300 ng/mL) for 8 h and lysed for immunoblotting for pERK1/2, ERK1/2, DNMT1 and EZH2. Representative image shown from 3 independent experiments. **(E)** OC cells (HeyA8 and Kuramochi) were treated with mesothelial cell conditioned medium (CM) with different inhibitors (AZD4547 or Volociximab) or with CM from mesothelial cells with IGFBP6 knocked down (IGFBP6 KD). RNA was isolated and qRT-PCR was performed for the expression of miR-193b-3p. All error bars represent mean ± SD; 3 independents experiments, p-value by Students t-test.

Once the unique secreted factors were identified, it was important to test their effect on activation of the ERK/EZH2/DNMT1 axis in the OC cells. Treatment with recombinant human bFGF and IGFBP6 caused an induction of ERK phosphorylation along with EZH2 and DNMT1 expression in a panel of OC cells (Figure 6D). Knocking down IGFBP6 in mesothelial cells made their conditioned medium incapable of inducing miR-193b-3p downregulation in OC cells (Figure 6E). Similarly, inhibiting FGFR prevented miR-193b-3p downregulation in mesothelial cell conditioned medium treated OC cells (Figure 6E). IGFBPs can inhibit IGFs but IGFs were not detected in the secretome arrays of mesothelial or OC cells (Figure 6A). IGFBPs can act independent of IGFs and induce cellular signaling through α5 and β1 integrins^22–24^. Therefore, the effect of inhibition of α5β1 integrin using a neutralizing antibody (Volociximab) was tested in OC cells treated with mesothelial cell conditioned medium. Volociximab treatment prevented the downregulation of miR-193b-3p by the conditioned medium (Figure 6E), confirming the role of α5β1 integrin as the IGFBP6 receptor.

### miR-193b-3p replacement therapy in an OC patient derived xenograft (PDX) model

Having identified the microenvironment-induced downregulation of miR-193b-3p being critical for initiation of OC metastatic colonization, the potential for miR-193b-3p replacement therapy was tested. A chemoresistant OC PDX model - AM, derived from patient ascites^25^ was selected for an intervention study involving treatment of established metastases with pre-miR-193b-3p or scrambled control oligos. We started by first testing if the patient ascites derived AM cells behaved similarly to OC cell lines upon interaction with the outer layers of the omentum. The AM OC cells were seeded on the 3D omentum culture model, sorted after 2 days of culture, and analyzed for the expression of miR-193b-3p, its targets cyclin and its upstream regulators. As observed with OC cell lines, miR-193b-3p was downregulated in the patient ascites derived AM cells, when seeded on the 3D omentum culture. Similarly, miR-193b-3p targets cyclin D1 and uPA, and its regulators EZH2 and DNMT1, were upregulated in AM cells seeded on the 3D omentum culture (Figure 7A and B). Moreover, analysis of a panel of OC stem cell markers confirmed that the omental microenvironment induced an OC stem cell phenotype in AM cells (Figure 7C). The activation of the upstream regulators of miR-193b-3p in AM cells was confirmed by western blotting (Figure 7D). Having confirmed that the effects observed in OC cell lines was also applicable to patient ascites derived OC cells, we proceeded to generate OC PDX by injecting the AM cells intra peritoneally (i.p.) in female NSG mice. Once metastases were established, mice were injected with pre-miR-193b-3p packaged with invivofectamine 3.0, twice a week. Treatment with miR-193b-3p resulted in significantly decreased metastases in the PDX compared to scrambled control oligo treatment (Figure 7E and F). This demonstrated the potential of miR-193b-3p replacement therapy to treat OC metastases.

**Figure 7:**
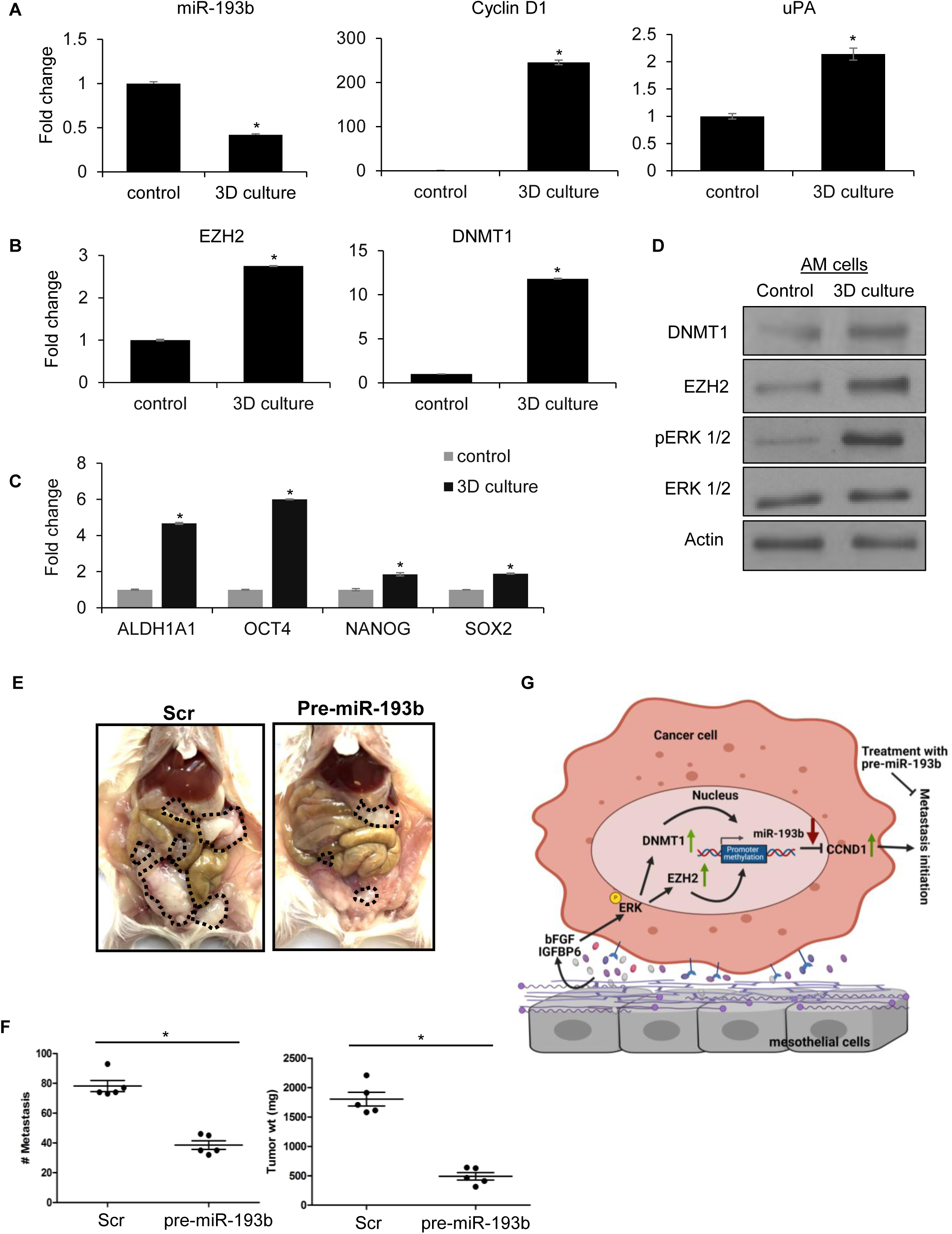
Validation in patient-derived tumor cells: AM cells were isolated from the ascites of an advanced-stage HGSOC patient. **(A)** CMFDA-labeled AM cells were seeded on the 3D omentum culture, allowed to grow for 48 h, and sorted by FACS. RNA was isolated, and qRT-PCR was performed for miR-193b-3p (*left*) and it targets CCND1 (*middle*) and uPA (*left*). Similarly, qRT-PCR was performed for EZH2 and DNMT1 **(B)** and for OC stem cell markers (ALDH1A1, OCT4, NANOG and SOX2) **(C)** (mean ± SD; 3 independents experiments). * p<0.01, Students t-test. **(D)** CMFDA-labeled AM cells were seeded on the 3D omentum culture, allowed to incubate for 48 h, and then isolated by FACS. Sorted cells were lysed for protein using RIPA buffer and immunoblotted for EZH2, DNMT1, phosphorylated ERK1/2 and ERK1/2. Representative image shown from 3 independent experiments. **(E)** miR-193b-3p replacement therapy: AM cells were injected i.p. into 6-week-old female NSG mice to establish metastases and then treated with 1mg/kg miR-193b-3p or scrambled negative control delivered in Invivofectamine 3.0 (200µL), injected i.p. on day 8, 13 and 18. The mice were euthanized on day 20 and the peritoneal metastasis were counted, surgically resected, and weighed. **(E)** Representative images of the peritoneal metastasis in each group with the tumors outlined in black. **(F)** The number of metastasis (*left*) and total tumor weight *(right)* in mice treated with pre-miR-193b-3p compared to scrambled control were plotted (mean ± SD; n=5 mice/group; *p<0.01). **(G)** Schematic overview of the findings: bFGF and IGFBP6 from mesothelial cells and CAFs in the microenvironment activate the ERK/EZH2/DNMT1 axis. EZH2 induces H3K7me3 at the miR-193b-3p promoter, which recruits DNMT1, catalyzing promoter hypermethylation and downregulation of miR-193b-3p. As a result, its target, cyclin D1 (stemness) is upregulated and imparts a MIC phenotype along with its other target uPA (invasiveness).

## Discussion

Recent research has demonstrated that a subset of cancer cells, which in addition to having cancer stem cell-like properties, have the ability to undergo EMT, survive stresses, and establish productive interactions with the metastatic microenvironment. This subset of cells is termed metastasis initiating cells (MICs) and have been reported in multiple malignancies^3^. Since the escape from the primary tumor during OC metastasis is relatively passive, involving exfoliation of the cancer cells into the peritoneal fluid, OC metastasis initiating cell studies are more relevant for their role in metastatic colonization. Successful metastatic initiation hinges on the intricate coordination of multiple cell types to build a microenvironment (niche) suitable for MICs to proliferate, differentiate, and populate the metastatic lesion.

We have identified a MIC suppressing microRNA, miR-193b-3p, which is downregulated in metastasizing OC cells by IGFBP6 and basic FGF secreted by mesothelial cells, via the ERK/EZH2/DNMT1 axis, promoting the MIC phenotype through upregulation of its targets Cyclin D1 (stemness) and uPA (invasion/growth) (Figure 7G). miR-193b-3p has been shown to be downregulated by promoter hypermethylation in prostate cancer and liposarcoma^26–28^. We have previously demonstrated that the metastatic microenvironment induces promoter hypermethylation and downregulation of miR-193b-3p in metastasizing OC cells^5^. Interestingly, the analysis of pan-cancer data from TCGA demonstrated that miR-193b-3p expression in primary tumors in OC is among the highest (Suppl Figure 7). Since escape from the primary tumor is passive, its downregulation during metastatic colonization is functionally relevant in the context of MICs. Here we show that the microenvironmental signaling downregulation miR-193b-3p involves a paracrine mechanism. FGF2 and IGFBP6 secreted by mesothelial cells covering the omentum activated ERK in the cancer cells during early metastatic colonization. In advanced metastases, this role in taken over by the CAFs. Interestingly, miR-193b-3p has been reported to regulate KRAS and thus ERK signaling in pancreatic cancer^29,30^. Therefore, our findings indicate the involvement of a potential feed forward mechanism for the ERK/miR-193b-3p axis. ERK induced EZH2, which was recruited to the miR193b-3p promoter and catalyzed H3K27^me3^. The regulation of EZH2 expression has also been previously demonstrated in lung cancers with KRAS mutations and in triple-negative and ERBB2-overexpressing subtypes of breast cancer^15,16^. DNMT1 was also induced by ERK activation and was recruited to the miR-193b-3p promoter by the increased H3K27me3, catalyzing promoter hypermethylation and downregulation of miR-193b-3p. The regulation of miR-193b-3p by DNMT1 has also been reported in liposarcoma^28^. Therefore, this mechanism is not limited to epithelial cancers.

The downregulation of miR-193b-3p induced an MIC phenotype of stemness and increased adhesion to the omentum coupled with increased invasiveness. CyclinD1 and urokinase were identified as the direct targets of miR-193b-3p with the former contributing to cancer stemness while the latter increased the invasiveness and motility of these cells. Cyclin D1 has been demonstrated as a direct target of miR-193b-3p in multiple malignancies^31–33^. Our data show its critical contribution towards the MIC phenotype in OC. A miR-193b-3p replacement therapy significantly reduced metastasis in a chemoresistant PDX. Exosomal delivery of miR-193b-3p has also been shown to be effective in inhibiting inflammatory response and neurodegeneration induced by subarachnoid hemorrhage in mice^34^. Similarly, nanoparticles packaged with miR-193b mimic have been tested in vivo in acute myeloid leukemias^35^. microRNA replacement therapies are being tested in clinical trials^36^. The most notable being MRX34, a liposomal miR-34a mimic, which was well tolerated and showed some antitumor activity in a subset of patients with refractory advanced solid tumors^37^. Taken together, our data indicate the potential for translating miR-193b-3p replacement therapy into a strategy to target MICs, thereby offering a promising approach to effectively treat OC patients, the majority of whom present with extensive metastasis.

## METHODS

### Reagents

All the OC cells were grown in Dulbecco’s Modified Eagle Medium (DMEM) (Corning), 10% FBS (Atlanta), 1% MEM vitamins (Corning), 1% MEM nonessential amino acids (Corning), and 1% Penicillin-Streptomycin solution (100x, Corning). TaqMan miRNA assay for hsa-miR-193b-3p was purchased from Applied Biosystems (Foster City, CA, USA). pre-miR-193b-3p mimic and scrambled or negative control were purchased from Ambion (Thermo Fisher Scientific, Hanover Park, IL). miRCURY LNA anti-miR-193 and scrambled anti-miR-negative control were from Exiqon (Vedbæk, Denmark). All the siRNAs used in the study were purchased from Dharmacon (Lafayette, CO, USA) SMARTpool siRNAs: Scrambled siRNA (Cat# D-001810-10-20), uPA siRNA (s10612), CCND1 siRNA (J-003210-16-0005). CCND1 overexpression plasmid vector (pcDNA3-Myc-CyclinD1(WT) addgene). Thiazolyl blue tetrazolium bromide (MTT reagent, Cat# 298-93-1) was from Acros Organics (Thermo Fisher Scientific, Hanover Park, IL) and 4% paraformaldehyde solution (Cat# NC9245948) was procured from Thermo Fisher Scientific (Hanover Park, IL).

### Cell lines

Human high-grade serous OC cell lines OVCAR3 and OVCAR8 were acquired from Ernst Lengyel (University of Chicago), OVCAR4 was from Joanna Burdette (University of Illinois at Chicago) and Kuramochi from Japanese Collection of Research Bioresources. The cell lines used were genetically validated and tested to be mycoplasma free using respective services from Idexx BioResearch (Columbia, MO). Patient derived primary cells (AM) were collected from the ascites of a patient with advanced-stage and “platinum-resistant” HGSOC and maintained as primary cultures in ultra-low attachment dishes in RPMI 1640 supplemented with 1% MEM-NEAA (Life technologies, USA), 2% B-27 (Gibco, USA), 1% insulin-transferrin-selenium (Gibco, USA), and 1% penicillin/ streptomycin (Corning, USA).

### Isolation and culture of primary cells (NOF and HPMC)

As described previously, human primary mesothelial cells (HPMC) and normal omental fibroblasts (NOF) were isolated from omentum from female donors ^38,39^. CAFs were isolated from OC patient metastasis as described previously ^40^.

### 3D culture setup

As described previously ^38,39^, the 3D omental culture was assembled in 10 cm culture dishes by first seeding (3.6 x10^5^) normal fibroblasts with collagen Type 1 (Rat tail, BD Cat#354236) in DMEM. After 6 h, HPMCs (3.6 x10^6^) were seeded on top of NOF embedded basement membrane to form a confluent monolayer resembling the mesothelium. The 3D culture was incubated for 24h to allow for secretion of factors, generating a complex microenvironment resembling the outer layers of the omentum. GFP-expressing OC cells (1 x10^6^) were seeded on the 3D culture and allowed to grow for 48h to mimic the attachment and initial events of colonization. Cells were trypsinized and GFP-labeled OC cells were isolated by fluorescence activated cell sorting (FACS) using BD FACS Aria II flow cytometer (BD Biosciences, San Jose, CA). Control cells were grown in regular culture dishes and underwent the same treatment as the 3D culture cells. RNA was isolated from the sorted OC cells using miRNeasy mini kit (Qiagen, Germantown, MD) using the manufacturer’s instructions.

### Coculture experiments

The HPMCs (6 × 10^6^) were seeded in 100 mm dishes, incubated for 18-24 h. GFP-tagged OC cells (1.5 - 2 × 10^6^) were overlaid onto the HPMCs and allowed to grow for 48 h to enable interactions. Cells were washed, trypsinized and fluorescent cancer cells were isolated by fluorescence activated cell sorting (BD FACS Aria II) and used for RNA isolation (miR-193b-3p and its targets) ^5^.

### Proximal culture

The proximal culture experiments were set up as described previously ^12^. OC cells were (300K/insert) were seeded in lower surface of a transwell insert with 0.4μm pores (Corning, Cat# # 3412) and allowed to incubate for 6 h for cells to attach. The insert was then flipped and placed in a well of a 6-well plate containing growth medium. The HPMCs were seeded (150K/insert) on the upper surface of the insert and the cells were allowed to grow for 48hrs. At the end point, cells were carefully trypsinized sequentially from each surface without intermixing and collected for RNA isolation and qRT-PCR.

### Transient transfection

OC cells were transiently transfected with 30 nM scrambled or control siRNA and pre-miR-193b-3p/anti-miR-193b-3p using TransIT-X2 (Mirus cat# MIR 6004) according to the manufacturer’s protocol. For plasmid transfections, 3 µg plasmid was transfected using TransIT-X2 as per the manufacturer’s protocol. The cells were used for experiments 48 h after transfection or as indicated ^39^.

### Real-time PCR

RNA was isolated from the cells using miRNeasy Kit (Qiagen Cat# 217004) and the RNA was used for reverse transcription using Applied Biosystems High-Capacity Reverse Transcription Kit (Cat# 4368813) for gene expression and Applied Biosystems® TaqMan® MicroRNA Reverse Transcription Kit (Cat# 4366597) for microRNA expression. qRT-PCR was performed for miR-193b-3p using Taqman miRNA assay (Assay ID 002367) according to the manufacturer’s protocol using a Roche LightCycler 96 system with U6 (001973) as endogenous control. Similarly, CCND1 (Hs00765553_m1) and uPA (Hs01547054_m1), DNMT1 (Hs00945875_m1), ALDH1A1 (Hs00946916_m1), OCT-4 (Hs00999634_gh), NANOG (Hs02387400_g1), SOX-2 (Hs01053049_S1), gene expression qRT-PCR analysis was performed using TaqMan gene expression assays according to the manufacturer’s protocol using Roche LightCycler 96 system. GAPDH (Hs00183740_m1) was used as an endogenous control. The expression of relative fold change was calculated using the ΔΔCt method ^39^.

### ALDH enzyme assay by imaging

As described previously^10^, ALDH enzymatic activity was measured using ALDEFLUOR assay kit (STEMCELL Technologies) according to manufacturer’s instructions. Briefly, the experimental OC cells were seeded in 24-well plate (Falcon, Cat# 353047) and allowed to incubate for 24 h. Incubate the cells with active ALDFLUORE reagent (5 µL/mL) with ALDEFLUOR assay buffer for 60 – 30 min depending on cell line and imaging the ALDH+ve cells using EVOS FL Auto microscope (Life Technologies). At least 5 different images were taken from 3 different technical replicates for each experimental condition. (mean ± SD; 3 independent experiments).

### ALDH enzyme assay by quantification

As described previously^10^, ALDH enzymatic activity was measured using ALDEFLUOR assay kit (STEMCELL Technologies) according to manufacturer’s instructions. Briefly, the experimental OC cells were trypsinized and collected in a 15-mL tube and the cells were incubated with active ALDFLUORE reagent (5 µL/mL) with ALDEFLUOR assay buffer for 60 – 30 min depending on cell line and fluorescence of the cells were measured by using LSRII flow cytometer (BD Biosciences) compared with cells incubated with DEAB control and counted by the quantification of the average three independent replicates for each experiment.

### Spheroid formation

As described previously^10^, OC cells were trypsinized and seeded in ultra-low attachment 24-well plate plates in stem cell media for spheroid formation assay. 1000-2000 cells were seeded in each well and cultured for 4-10 days and imaging using EVOS FL Auto Imaging System (Life Technologies). At least 3 different images were taken from 3 different replicates. Spheroids were manually quantified. (mean ± SD; 3 independent experiments).

### Sorting the ALDH +ve and –ve cells

As described previously^10^, OC cells (OVCAR3) were trypsinized and mixed with (5 µL/mL) activated ALDEFLUOR reagent according to the manufacturer’s protocol (Stem Cell Technologies; Catalog #01700), along with DEAB control. Cells were incubated at 37 ^0^C, 5% CO_2_ incubator for 45 min. Cancer cells that express stem cell marker enzyme aldehyde dehydrogenase (ALDH), were separated by fluorescence activated cell sorting (BD FACS Aria II). Collected ALDH +Ve and -Ve cells were grown in DMEM medium, after 48 h cells were labeled with CMFDA and allowed to incubate for 60-90 min for recovery.

### In vivo adhesion

ALDH^+^ or ALDH^-^ OVCAR3 cells were sorted as described above. NSG mice (3 per group) were injected with CMFDA labeled ALDH^+^ or ALDH^-^ (350K/0.5 mL/mouse). After 3 hours, mice were euthanized to collect their peritoneum and omentum. The tissues were washed with PBS three times to remove the non-adherent cells and blood. Washed tissues were transferred into 6-well plate containing diluted trypsin with PBS (1:1) and allowed to incubate for 20 min with gentle shaking. Trypsinized cell suspension was collected, centrifuged, and the pellet was resuspended in PBS to measure the fluorescence using a SynergyH1 plate reader (Agilent Technologies).

### 3D Invasion

The 3D omental culture was assembled as described previously^5,6,39^, in FluoroBlok transwell inserts with 8 μm pores. Briefly, (4×10^3^) normal fibroblasts mixed with collagen Type 1 (Rat tail, BD Cat#354236) were seeded in FluoroBlok transwell inserts (8um pores) in DMEM and incubated at 37^0^C for 5 hours. Thereafter, a confluent monolayer of HPMCs were seeded on top of this basement membrane and allowed to attach overnight. CMFDA labeled OVCAR3 (7.5×10^4^/500 µl) cells (ALDH +Ve and -Ve) were seeded in serum free medium on the 3D culture and allowed to invade. 700 µl of DMEM with 10% FBS served as the chemoattractant in the lower chamber. Invasion is stopped after 16 h by fixing with 4%PFA for 20min at room temperature, washed with PBS and the invaded fluorescent OC cells imaged (5 fields/insert) using the EVOS FL auto microscope (Life Technologies).

### Chromatin immunoprecipitation (ChIP)

ChIP assay was performed as described by manufacturer protocol, briefly, HeyA8 cells were treated with U0126 (10 µM/mL) or DMSO for everyday (3 days) and harvested on 4^th^ day. Cells were cross-linked in a 1% formaldehyde solution and followed the ChIP-IT® Express Chromatin Immunoprecipitation Kits (ActiveMotif, Cat#53008). Decross-linked cells were sonicated into fragments (500-bp), using a Diagenode Bioruptor Pico (Diagenode, Denville, NJ, USA). Magnetic Dyna beads (ActiveMotif), combined with a mixture of specific antibodies DNMT1 (Epigenetek, Cat# A-1700), EZH2 (Cell signaling Technologies, Cat#5246) and K327me3 (Diagenode, Cat# C15410069) along with IgG as control. Immunoprecipitated DNA-protein complexes cross-links were reversed, and the DNA mixture then digested with proteinase K. The pulldown DNA was washed and purified by QIAquick PCR purification kit (Qiagen, Cat#28104). DNA fragments were recovered and used as templates for qPCR amplification (FastStart Essential DNA Green Master kit, Roche, Cat#06402712001), using specific primers for the miR-193b-3p promoter (GAAACUCCCGGUCA).

### Methylated DNA Immunoprecipitation (MeDIP)

Methylated DNA Immunoprecipitation was done as described by manufacture protocol (Active Motif Cat#55009), briefly, HeyA8 cells were treated with U0126 (10 µM/mL) or DMSO for everyday (3 days) and harvested on 4^th^ day. The cells were harvested and isolate the DNA using (DNeasy Blood & Tissue kit, Qiagen, cat#69504). Then the DNA was fragmented using sonication and fragmented DNA was mixed with Buffer C, to convert the single strand DNA. The fragmented DNA was Immunoprecipitated with 5-methylcytocine (5-mc) antibody or mouse IgG (control) along with bridging antibody for overnight at 4^0^C. Wash and elute the methylated DNA using QIAamp DNA micro-Kit (Cat#56304). Methylated DNA was further validated for enrichment in the miR-193b-3p promoter region using syber green qPCR.

### Secretome Array

HGF array C1 (Raybiotech, Cat#AAH-GF-1-2) was used to quantitatively compare the differences in secreted growth factors between mesothelial cells (n=3) and OC cells (OVCAR3, OVCAR4, Kuramochi) in accordance with the manufacturer’s protocol.

### Immunoblotting

Immunoblotting was done as previously described ^39^. Briefly, proteins were separated by 4-20% gradient SDS-PAGE and transferred to nitrocellulose membrane, probed with phospho-ERK 1/2 (Cell Signaling, Cat# #9101), ERK 1/2 (Cell Signaling, Cat# #4695), DNMT1 (Epigenetek, Cat# A-1700), EZH2 (Cell signaling, Cat#5246) K3H27Me3 (Diagenode, Cat# C15410069), uPA (Sekisui diagnostics, Cat#ADG3689), CCND1 (Cell signaling, Cat#2978), Actin (Sigma Aldrich, Cat# G9295) was probed as a loading control. Horseradish peroxidase-linked anti-rabbit (Cell Signaling, Cat# 7074) or anti-mouse (Cell Signaling, Cat# 7076) IgG secondary antibody was used to probe the membrane bound primary antibodies and detected using clarity western ECL substrate (BioRad, Cat# 170-5061).

### Immunofluorescence

As described previously^39^ the 3D omental culture was assembled on polyD-lysine coated glass coverslips and OC cells expressing GFP or RFP were seeded on the 3D culture or on the cover slips. Cells were fixed at 48h with 4% paraformaldehyde and permeabilized, blocked, and probed with primary antibodies against ERK1/2 (Cell Signaling, Cat #4695), phosphor-ERK1/2 (Cell Signaling, Cat# #9101), DNMT1 (Epigenetek, Cat# A-1700) or EZH2 (Cell signaling, Cat#5246) and then with Alexa Fluor 594 conjugated secondary antibody (Cell Signaling, Cat#8889S). Nuclei were stained with Hoechst 33342 (ThermoFisher Cat# H3570) and the coverslips were mounted with ProLong Gold (ThermoFisher Cat# P36934) and imaged using a Leica SP8 confocal microscope.

### Mesothelial cell conditioned medium

Mesothelial cells (6×10^6^) were seeded in a 10 cm dish and allowed to attach overnight. Thereafter, medium was aspirated, cells were washed 3 ties with PBS and 10 ml of serum free medium was added. The conditioned medium was collected after 24 h and concentrated using Amicon Ultra-15 centrifugal filters.

### RNA-seq

Next Seq reads were trimmed using Trimmomatic version 0.36 with the parameters “2:20:5 LEADING:3 TRAILING:3 SLIDINGWINDOW:4:15 MINLEN:17” to remove adapter sequences and perform quality trimming^41^. The resulting reads were mapped against GRCh38 using HISAT2 version 2.1.0 with default parameters^42^. HISAT uses Bowtie2, which is based on the Burrows-Wheeler transform algorithm, for sequence alignment and allows for mapping across exon junctions^43^. Read counts for each gene were created using feature Counts from the Subread package version 1.6.4 with the parameters “-O -M --primary --largest Overlap -s 2 -B” and Gencode v31 as the annotation^44,45^. Differential expression analysis was performed using the DESeq2 package (version 1.24.0) in R/Bioconductor (R version 3.6.0)^46^. The Venn diagram was also created in R using the Venn Diagram package (version 1.6.20).

### Preprocessing of scRNA-seq Data

The filtered gene expression count matrix was obtained from the publicly available dataset GSE165897 [PMID: 35196078]. To mitigate patient-specific batch effects, we employed the integration workflow implemented in Seurat v4.2.1^47–50^, following the recommended procedure described in the official tutorial (https://satijalab.org/seurat/articles/integration_introduction.html). Integration was performed using 8,000 selected features to generate a unified expression matrix. *Imputation of scRNA-seq Data:* To address the high dropout rates inherent in single-cell RNA sequencing data, we applied MAGIC (Markov Affinity-based Graph Imputation of Cells) v2.0.3^51^ for data imputation. The MAGIC algorithm was run with parameters k = 20 and t = 3 on the integrated expression matrix to recover gene expression signals and preserve the underlying biological structure.

#### Annotation of scRNA-seq Data

Cell clustering was done using a graph-based approach, utilizing the Louvain algorithm based on a shared nearest neighbor (SNN) graph^48^. To annotate different cell subpopulations, we utilized SingleR v1.10.0^52^, which identified all stromal cells as fibroblasts. To prevent misclassification of mesothelial and endothelial cells as fibroblasts, we further evaluated the expression of canonical markers, including CALB2 for mesothelial cells, and PECAM1 for endothelial cells. Fibroblast subpopulations were subsequently isolated from the stromal compartment and subjected to downstream analyses. To further refine the annotation of fibroblasts, endothelial cells, mesothelial cells, and cancer cells, we applied the same graph-based clustering algorithm to each respective cell subpopulation to identify distinct subclusters within each cell type.

#### Dimensionality Reduction and Visualization

To visualize the high-dimensional single-cell transcriptomic data, we performed dimensionality reduction using Uniform Manifold Approximation and Projection (UMAP)^53^ as implemented in Seurat v4.2.1. UMAP was applied to the top principal components (typically the first 30 PCs), which were identified via principal component analysis (PCA) on scaled expression values. This approach preserves both local and global transcriptomic structure, enabling effective visualization of cellular heterogeneity and clustering results in a two-dimensional space.

### Patient derived Xenograft Experiments

The mouse xenograft experiments were conducted following protocols approved by the Indiana University Bloomington Institutional Animal Care and Use Committee. Patient ascites derived OC cells (AM cells, 10 × 10^6^/ 500uL)^25,54^ were injected intraperitoneally into 6-week-old female NSG mice, mice were treated with 3 rounds of miR-193b-3p or scrambled negative control (1mg/kg) using Invivofectamine 3.0 (200µL) (thermos Fisher Scientific, Cat # IVF3001) and the mice were euthanized on 20th day and the peritoneal metastasis were counted, surgically resected, and weighed as described previously^5,6,39^.

### Statistics

Data analysis was done by unpaired, two-tailed Student’s t-test assuming equal variance of the test and the control populations.

## Acknowledgments

We are indebted to all the patients for their participation in tissue collection for these experiments. Funding sources: AKM was supported by a DoD OCRP Ovarian Cancer Academy Award (W81XWH-15-0253) and an American Cancer Society Research Scholar Grant (RSG-21-019-01-CSM).

