## Supplementary figures 1-7 for "Metastatic niche mediated activation of metastasis initiating cells in ovarian cancer through miR-193b-3p downregulation via the ERK/EZH2/DNMT1 axis"

**A**

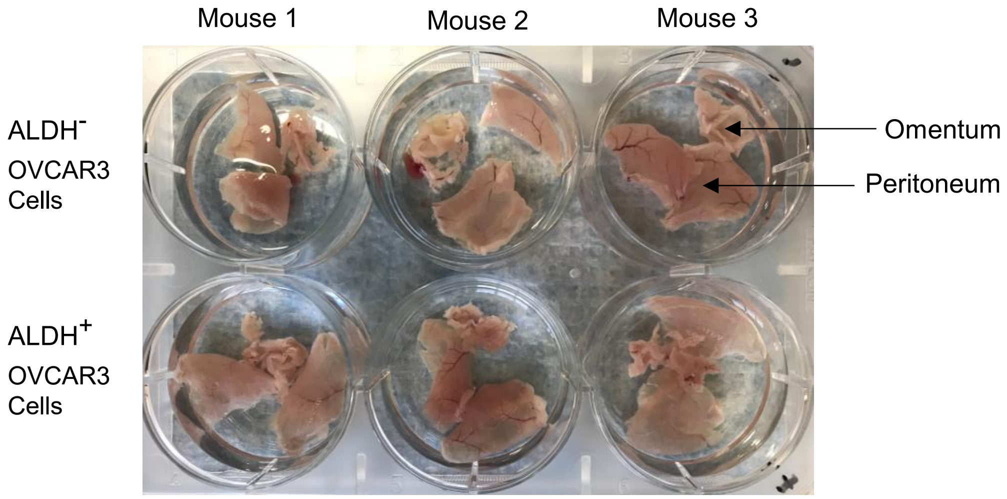


**B**

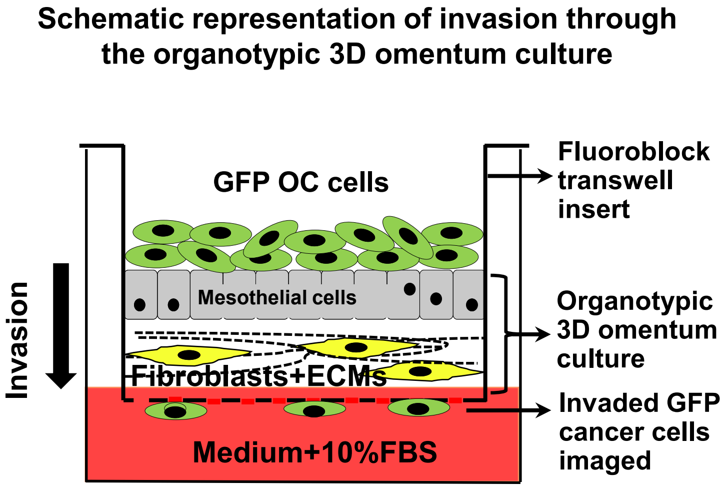


**Supplementary Figure 1: A)** In vivo adhesion assay: Images of mouse omentum and peritoneum removed to extract adherent fluorescent OC cells. **B)** Invasion through the outer layers of the omentum. The organotypic 3D omentum culture, mimicking the outer layers of the omentum, is assembled in fluoroblock transwell inserts. GFP OC cells are then seeded on the 3D omentum culture and allowed to invade through it. Medium with 10% FBS acts as a chemoattractant in the lower chamber. Invaded GFP OC cells are imaged from the bottom using EVOS FL fluorescent microscope and counted.

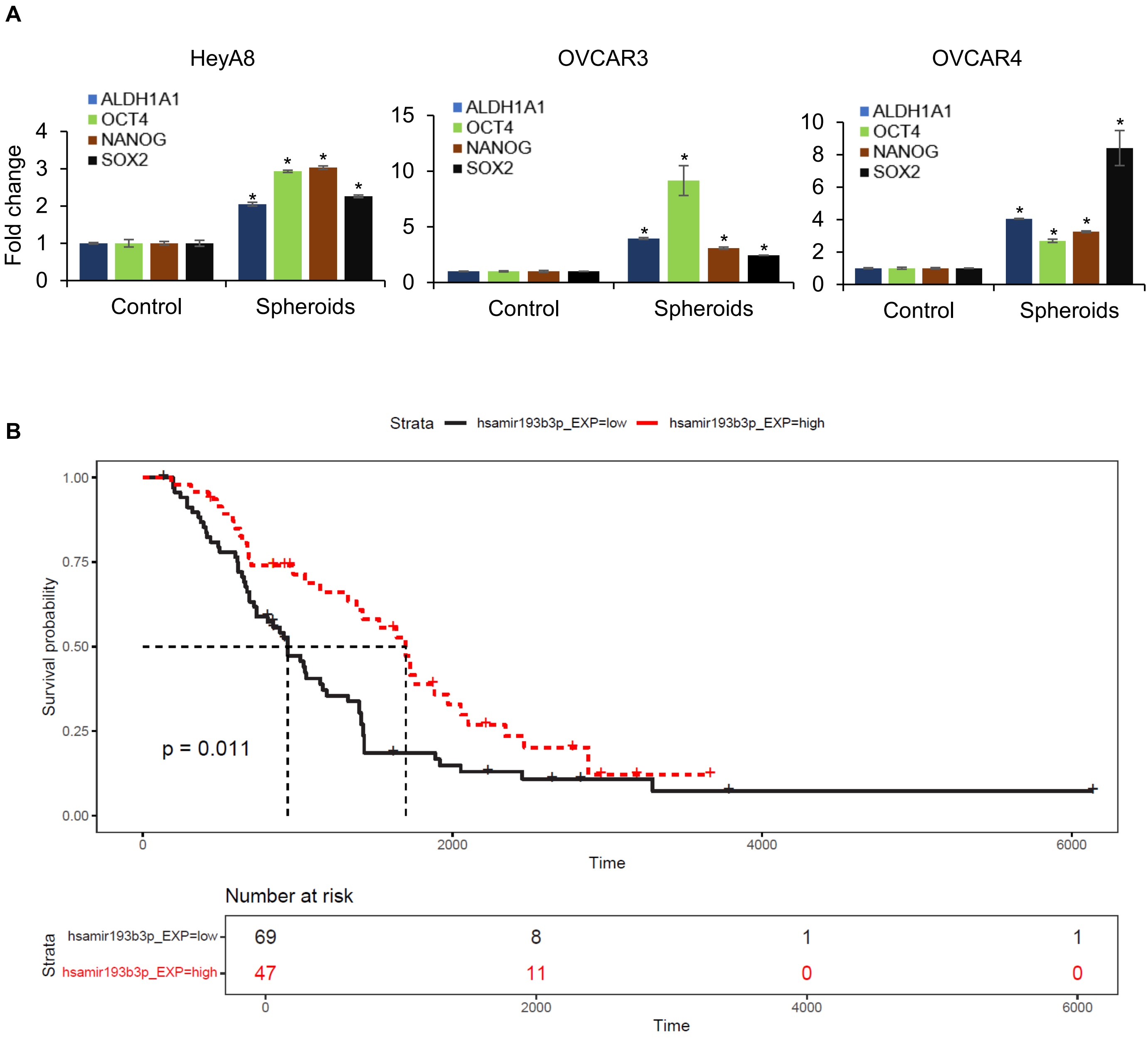


**Supplementary Figure 2: A)** OC cell spheroids were grown in an ultra-low attachment 6-well plate (HeyA8 – 3 days, OVCAR3 – 10 days and OVCAR4 – 7 days). RNA was isolated and qRT-PCR was done for stem cells markers (ALDH1A1, OCT4, NANOG and SOX2) by comparing the control cells (adherent cells growing in 2D) Mean ± SD from 3 independent experiments; * $p<0.01$. **B)** Kaplan-Meier survival curve for hsa-mir-193b-3p expression from Australian Ovarian Cancer dataset downloaded from the ICGC data portal. The black line represents patients with low (n=69), and red represents high expression (n=47). The log-rank test shows a statistically significant difference between the two groups (p = 0.011), with patients having low hsa-mir-193b-3p expression demonstrating shorter survival times compared to those with high expression. Cox proportional hazards regression analysis revealed a hazard ratio of 0.5726 (95% CI: [0.371 - 0.8836], p = 0.01177702) for high vs. low hsa-mir-193b-3p expression.

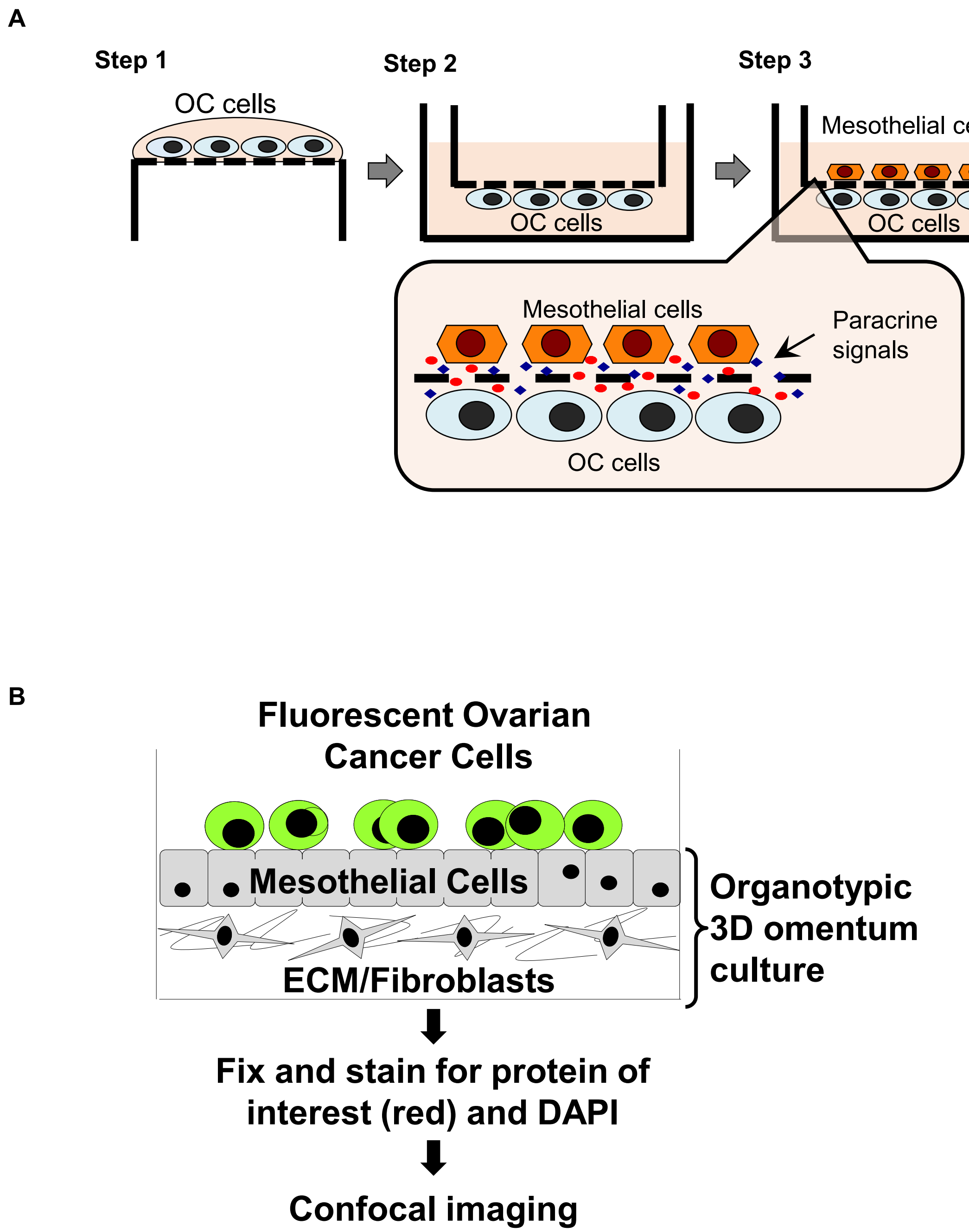


**Supplementary Figure 3: A)** Proximal culture schematic: OC cells were seeded on the bottom surface of a transwell insert with 0.4 μm pores. The insert was flipped and mesothelial cells were seeded on the opposite surface. Cells are then cultured, allowing exchange of secreted factors at localized high concentrations while preventing direct cellular contact. **B)** Schematic for immunofluorescent staining of proteins of interest in fluorescent OC cells seeded on the organotypic 3D omentum culture.

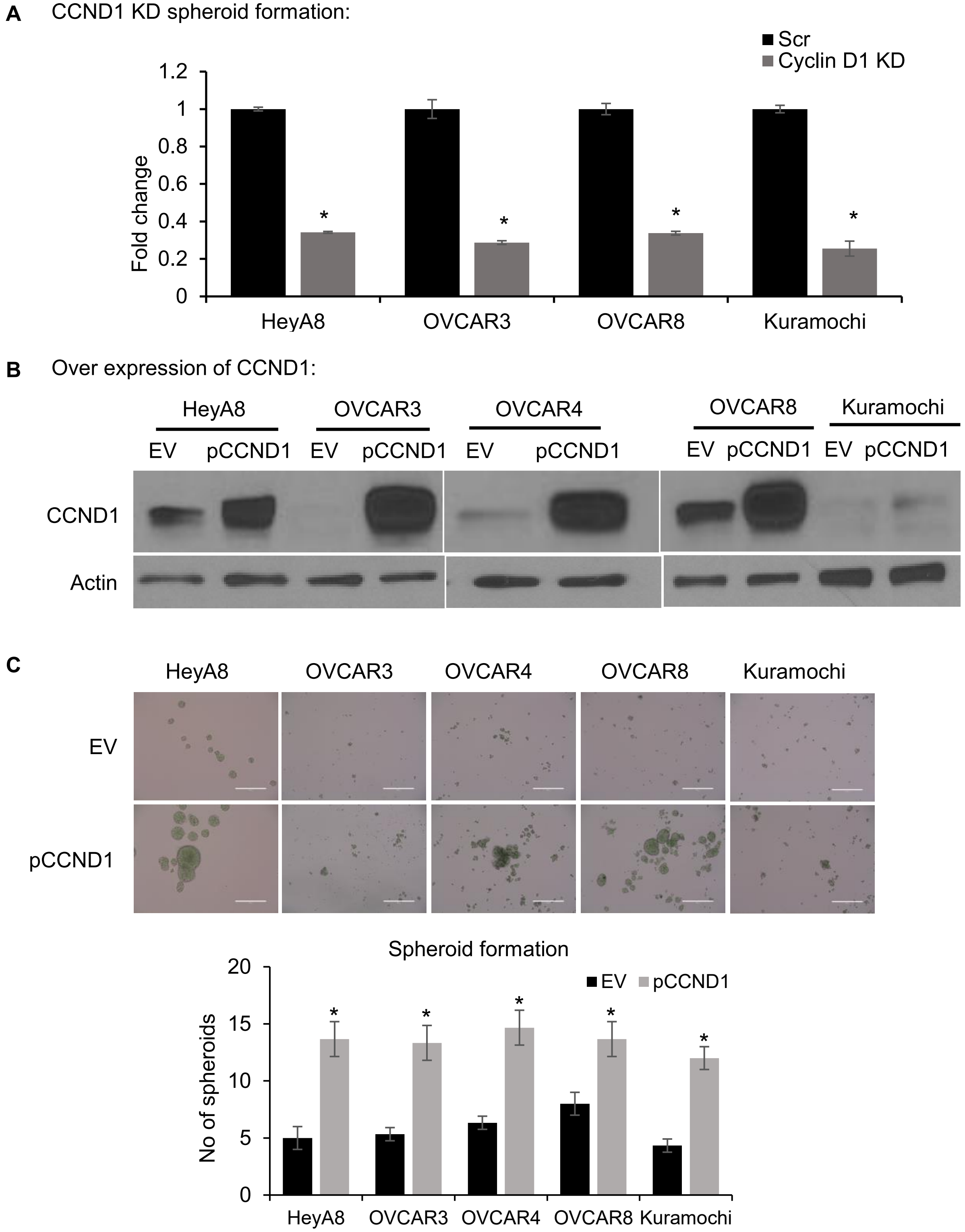


**Supplementary Figure 4: A)** Spheroid formation assay performed with OC cells transfected with CCND1 siRNA or scrambled control oligos (scr). Spheroids were imaged with an EVOS FL Auto microscope, counted, and plotted. **B)** OC cells (HeyA8, OVCAR3, OVCAR4, OVCAR8 and Kuramochi) were transfected with empty vector (EV) and CCND1 overexpression plasmid. Cells were lysed after 3 days for immunoblotting for the expression of CCND1. **C)** OC (HeyA8, OVCAR3, OVCAR4, OVCAR8 and Kuramochi) cells were transfected with EV an pCCND1 and were grown in ultra-low attachment plates for the spheroid formation. Spheroids were imaged with an EVOS FL microscope and quantified (EV – empty vector, pCCND1 – Cyclin D1 overexpression vector). All error bars represent mean ± SD; 3 independents experiments, * p<0.01, Students t-test.

**A** qPCR for miR-193b-3p

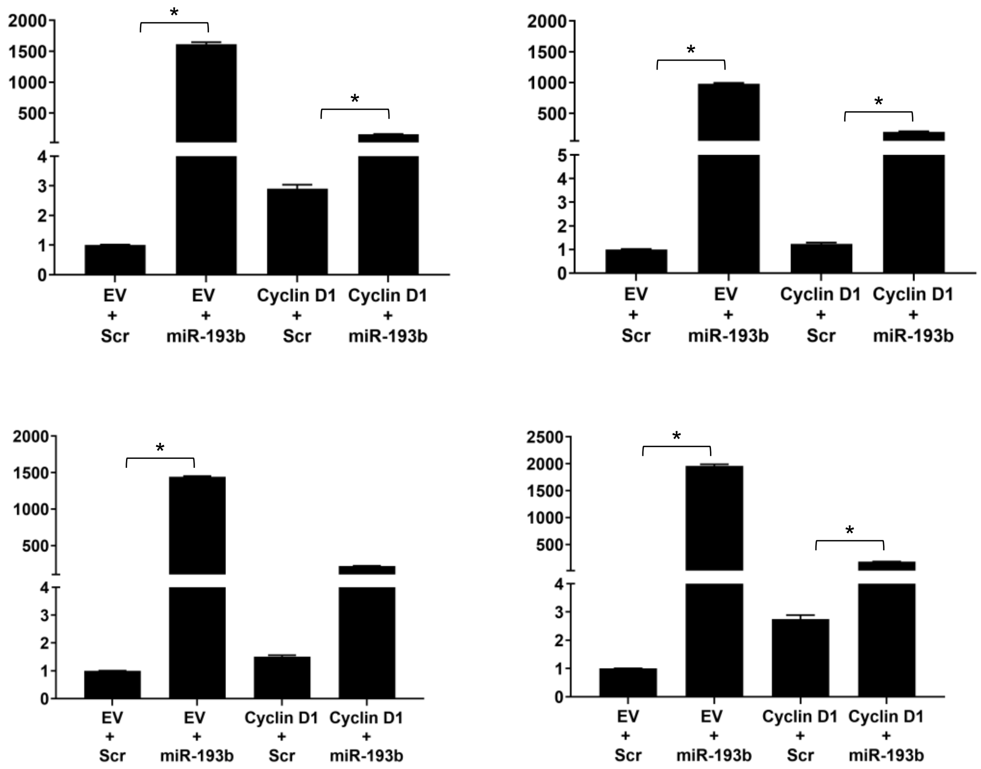


**B** WB for Cyclin D1

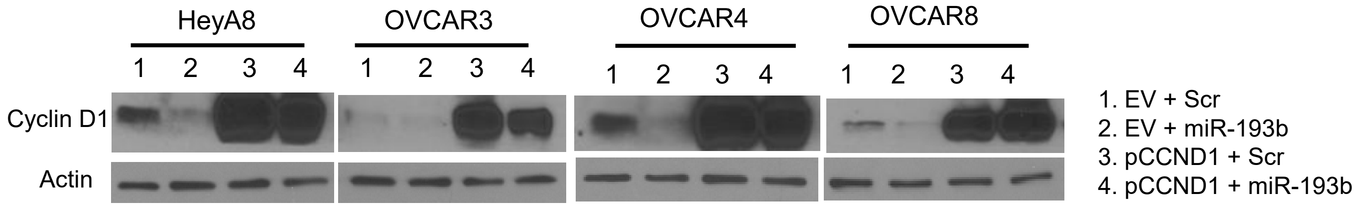


**Supplementary Figure 5: A)** OC cells were co-transfected with empty vector (EV)/cyclin D1 overexpression plasmid (CCND1) or scrambled negative control oligo (Scr)/pre-miR-193b-3p (miR-193b) as indicated. RNA was isolated after 2 days for qRT-PCR for miR-193b-3p. Error bars represent SD from 3 independent repeats, * p<0.01. **B)** OC cells were co-transfected with empty vector (EV)/cyclin D1 overexpression plasmid (CCND1) or scrambled negative control oligo (Scr)/pre-miR-193b-3p (miR-193b) as indicated and lysed after 3 days for immunoblotting for cyclin D1. Representative images from 3 independent repeats.

**A** Secretome array

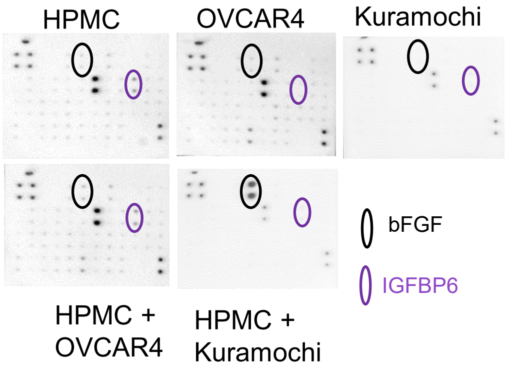


**B** Expression of bFGF and IGFBP6 mRNA

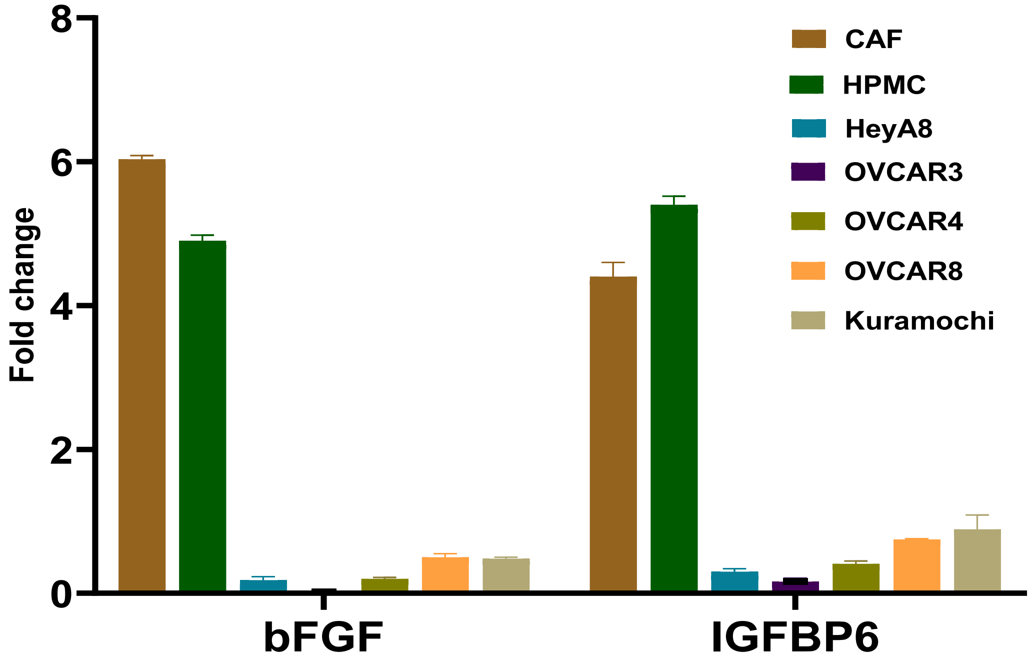


**Supplementary Figure 6: A)** Secretome array images of corresponding CM from human primary mesothelial cells (HPMC), OC cells (OVCAR4 and Kuramochi), and their coculture. **B)** Expression of bFGF and IGFBP6 mRNA in HPMCs and OC cells, RNA was isolated from HPMCs and OC cells, qRT-PCR was performed for bFGF2 and IGFBP6. Error bars represent standard deviation from 3 independent repeats.

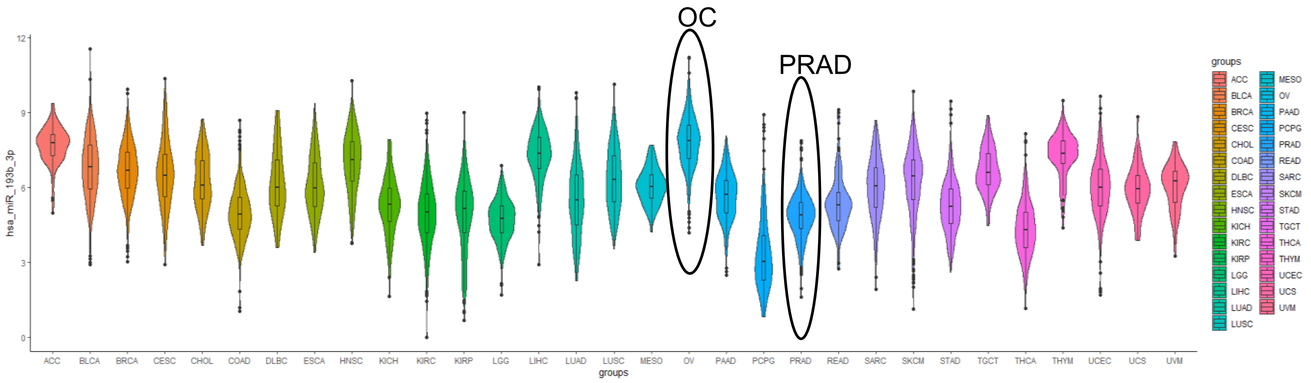


**Supplementary Figure 7: E**xpression of miR193b-3p in pan-cancer dataset from TCGA; OC: ovarian cancer; PRAD: Prostate adenocarcinoma
